# Neuronal activity remodels the F-actin based submembrane lattice in dendrites but not axons of hippocampal neurons

**DOI:** 10.1101/2020.05.27.119453

**Authors:** Flavie Lavoie-Cardinal, Anthony Bilodeau, Mado Lemieux, Marc-André Gardner, Theresa Wiesner, Gabrielle Laramée, Christian Gagné, Paul De Koninck

## Abstract

The nanoscale organization of the F-actin cytoskeleton in neurons comprises membrane-associated periodical rings, bundles, and longitudinal fibers. The F-actin rings have been observed predominantly in axons but only sporadically in dendrites, where fluorescence nanoscopy reveals various patterns of F-actin arranged in mixed patches. These complex dendritic F-actin patterns pose a challenge for investigating quantitatively their regulatory mechanisms. We developed here a weakly supervised deep learning segmentation approach of fluorescence nanoscopy images of F-actin in cultured hippocampal neurons. This approach enabled the quantitative assessment of F-actin remodeling, revealing the disappearance of the rings during neuronal activity in dendrites, but not in axons. The dendritic F-actin cytoskeleton of activated neurons remodeled into longitudinal fibers. We show that this activity-dependent remodeling involves Ca^2+^ and NMDA-dependent mechanisms. This highly dynamic restructuring of dendritic F-actin based submembrane lattice into longitudinal fibers may serve to support activity-dependent membrane remodeling, protein trafficking and neuronal plasticity.

## Introduction

One of the hallmark discoveries made possible by fluorescence nanoscopy methods is the existence of a periodical lattice of F-actin, spectrin, and associated proteins under the surface membrane of neuronal processes. This lattice, containing F-actin rings periodically spaced 180-190nm apart, was initially discovered in axons [1]. The lattice was later observed in dendrites of multiple types of neurons, albeit to a lesser extent compared to axons [2–4]. Several isoforms of spectrin have been observed in the lattice, with variable prevalence in axons compared to dendrites and during development [3, 4]. In addition to forming periodical rings, other F-actin structures have been described at the nanoscale in axons and dendritic shaft, including longitudinal fibers [2, 5–7].

The role and regulatory mechanisms of the periodical submembrane skeletal structure remain unclear. It has been shown to be regulated during development in axons, dendrites, and dendritic spines [2–4, 8]. It was demonstrated that the submembrane lattice destabilization triggers axonal degeneration [9, 10]. A recent study provided evidence that it serves as a signaling platform for receptor tyrosine kinase transactivation in neurons [11].

The more variable appearance and sporadic presence of the F-actin/spectrin lattice in dendrites compared to axons suggests that the structure is differently regulated in these distinct processes [2–4, 12]. A clear discrimination between spatially overlapping axons and dendrites, using a combination of specific markers, has been lacking in studies comparing the properties of the lattice, which may have impacted the analyses of its prevalence in dendrites. Furthermore, the greater diversity of F-actin nanostructures in dendrites compared to axons poses an additional challenge for quantitative analysis of the submembrane lattice. Several features of neuronal culture conditions, age, or fixation methods may have also contributed to discrepancies across studies of the dendritic lattice. One feature that has not been controlled or assessed is the level of electrical or synaptic activity. Ample evidence has shown, using conventional microscopy methods, that neuronal activity regulates the F-actin cytoskeleton [13, 14].

We thus set out to test whether neuronal activity regulates the F-actin-based lattice in dendrites and axons of cultured hippocampal neurons, using STimulated Emission Depletion (STED) nanoscopy. The complexity of the nanoscale F-actin patterns in dendrites and axons required the use of image segmentation to quantify their dynamics. The high diversity and variability of the reported F-actin patterns [1, 2, 4, 6, 8] prompted a high throughput analysis framework for the detection of those patterns on nanoscopy images. Recently, deep learning methods have been developed for automated feature detection in microscopy images from cells [15–19]. We show here that, using a weakly supervised deep learning approach [20–22], we could train a modified U-Net architecture [15] to segment regions containing fluorescent F-actin rings and/or longitudinal fibers in axons and dendrites. We demonstrate that this can be done without a significant decrease in segmentation quality, even in the presence of incomplete or coarse labeling. Our quantitative analysis highlights the profound diversity of F-actin patterns in dendrites, revealing an activity-dependent remodeling, from a lattice pattern to longitudinal fibers, which does not occur in axons, involving the influx of Ca^2+^. The activity-dependent remodeling of F-actin structures in dendrites may explain the previous sporadic observations of the submembrane F-actin lattice in dendrites and may be necessary to regulate membrane dynamics and protein transport required for dendritic signaling and plasticity.

## Results

### Activity-dependent remodelling of dendritic but not axonal F-actin nanostructures

To resolve the nanoscale organization of the F-actin cytoskeleton in neurons, we used the F-actin fluorescent marker phalloidin-STAR635, in low density cultured rat hippocampal neurons (8 and 13 DIV), imaged with STED nanoscopy. We observed complex and diverse patterns of fluorescence inside the neuronal processes that were only distinguishable with super-resolution microscopy (Fig. 1). The F-actin periodical ring patterns were robustly detectable in axons (co-labelled with axonal marker SMI31 antibody, detecting phosphorylated neurofilaments, enriched in axons [23])(Fig. 1). By contrast, dendrites (co-labelled with dendritic marker MAP2 antibody) exhibited patches of i) F-actin rings perpendicular to the shaft, mixed with patches of what appeared as either ii) unstructured patterns of F-actin, iii) compact assemblies of F-actin, or iv) longitudinal fibers parallel to the shaft axis (Fig. 1) [2, 6, 8]. We also observed a polygonal-like (2D) F-actin lattice in somatic regions, as previously described [4] (Supplementary Fig. 1a). These results confirm previous work illustrating the difference in the patterns of F-actin labelling in axons and dendrites [2–4]. They also highlight the frequent spatial overlap of axons and dendrites in cultures, revealing clear axonal lattice patterns overlaying dendrites (Fig. 1 orange arrowheads). Hence the triple staining (F-actin, dendrites, axons) is necessary to investigate the regulation of the lattice in both compartments.

**Figure 1:**
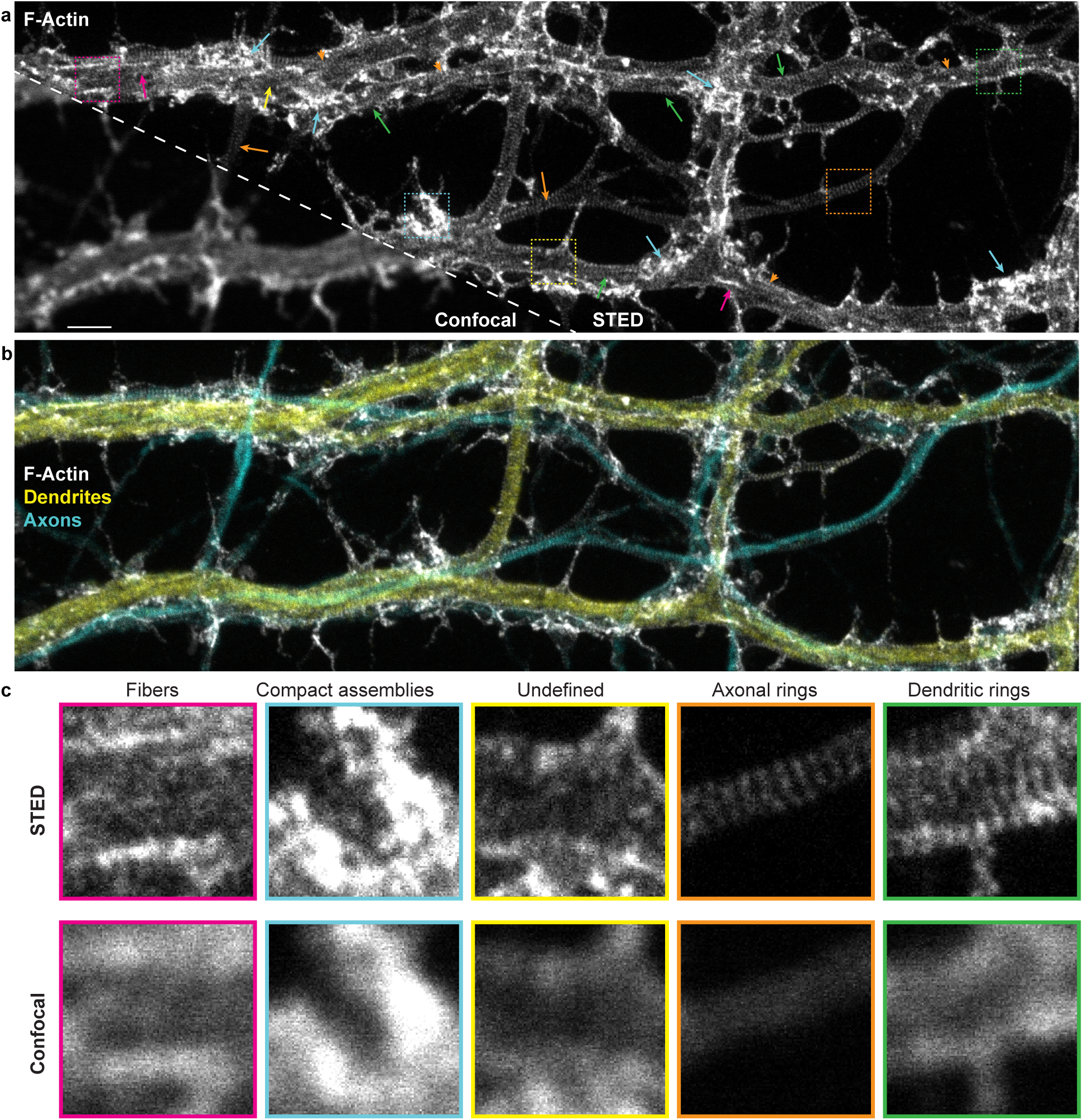
STED nanoscopy reveals diverse nanostructures of F-actin, in cultured hippocampal neuronal processes, that cannot be resolved with confocal microscopy. a) Representative image of the F-actin skeleton showing the diversity of nanostructures that can be observed with STED nanoscopy. Arrows point to regions exhibiting dendritic rings (green), axonal rings (orange), longitudinal fibers (magenta), compact assemblies (cyan), or undefined or diffuse signal (yellow). Orange arrowheads indicate regions of overlapping axonal and dendritic patterns. b) Three color imaging of the region in a) showing the overlap between axons (cyan, phosphorylated neurofilaments - SMI31) and dendrites (yellow, MAP2). MAP2 and SMI31 were imaged with confocal resolution to highlight the shape of the processes. c) Insets show a magnification of the regions indicated with the dashed squares in a) for both STED (top) and confocal (bottom) imaging modalities. Scale bar 2 *µ*m, insets 3.2 × 3.2*µ*m.

To investigate the dynamics of the submembrane lattice in axons and dendrites, via the monitoring of F-actin patterns, we considered a number of labeling strategies that were previously exploited for F-actin imaging at the nanoscale. These include photoactivatable green fluorescent protein (GFP), Lifeact, and Actin-Chromobody. However, these labels were shown not to reveal the F-actin periodical ring structure [24–26]. We also considered and tested live staining of the F-actin rings with fluorescent Jasplakinolide-derivative, a drug known to promote actin polymerization [2, 27, 28]. We observed that this reagent, tagged with silicone rhodamine (SiR-actin), revealed both F-actin rings and longitudinal fibers, as previously shown (Supplementary Fig. 2a-c) [2]. However, it was also shown that the reagent altered F-actin dynamics [29]. In fact, we noticed that the presence of F-actin fibers and bundles increased with incubation time, higher concentration of the SiR-actin label, or even post-incubation delay. To measure the impact of this Jasplakinolide-derivative on the presence of F-actin rings and longitudinal fibers, we incubated the neurons with a very low concentration of SiR-actin (0.5 *µ*M) for only 8 min, well below the manufacturer specification. We then fixed the cells with PFA and stained with phalloidin-STAR635 to quantify the F-actin patterns. The results show that SiR-actin pre-incubation increases the prevalence of F-actin longitudinal fiber over rings in dendrites compared to untreated neurons. These results confirm that the SiR-actin interferes with F-actin dynamics in live neurons, indicating that this approach is unsuitable to investigate the activity-dependent dynamics of F-actin remodeling. (Supplementary Fig. 2d).

We thus decided to fix the neurons and triple stain them with phalloidin-STAR635 along with dendritic and axonal markers (Fig. 1, 2, and Supplementary Fig. 3) to further assess the F-actin regulation in both neuronal compartments. We next asked whether neuronal activity had an impact on the patterns of F-actin in axons and dendrites, by applying activity-promoting/inhibiting treatments on live neurons, which we then fixed and triple-stained. We manipulated neuronal activity by incubating coverslips of neurons in either i) a high Mg^2+^/low Ca^2+^ solution (10 min) to reduce basal neuronal activity, ii) a 0Mg^2+^/glycine/bicuculline (0Mg^2+^/Gly/Bic) solution (10 min) to promote excitatory synaptic and NMDA receptor activity, iii) a high K^+^ (40 mM) solution (2 min) to briefly depolarize neurons, or iv) a glutamate/glycine (Glu/Gly) solution (2 min) to produce a broad excitatory stimulation [30, 31]. Using STED nanoscopy, we observed activity-dependent remodelling of F-actin nanostructures on dendrites that could not be resolved with diffraction-limited confocal microscopy (Fig. 2). Increasing neuronal activity led to the reorganization of F-actin from periodical rings to longitudinal fibers (Fig. 2, green and magenta arrows). While in the low activity, high Mg^2+^/low Ca^2+^ condition, dendritic F-actin rings were prevalent, the strong activity promoting stimulation Glu/Gly resulted in F-actin longitudinal fibers being predominant on most of the dendritic shaft. The brief High K+ treatment induced a less pronounced reorganisation, while the synaptic stimulation 0Mg^2+^/Gly/Bic was associated with a patchy F-actin lattice of intercalating or overlapping rings and fibers. Strikingly, we observed little impact of any of these treatments on the presence of the axonal periodical F-actin ring pattern (Fig. 2, orange arrows). These results suggest that while the submembrane periodical F-actin rings in axons are unaffected by neuronal activity, the dendritic rings disappear with increasing neuronal activity, seemingly reorganizing into longitudinal F-actin fibers.

**Figure 2:**
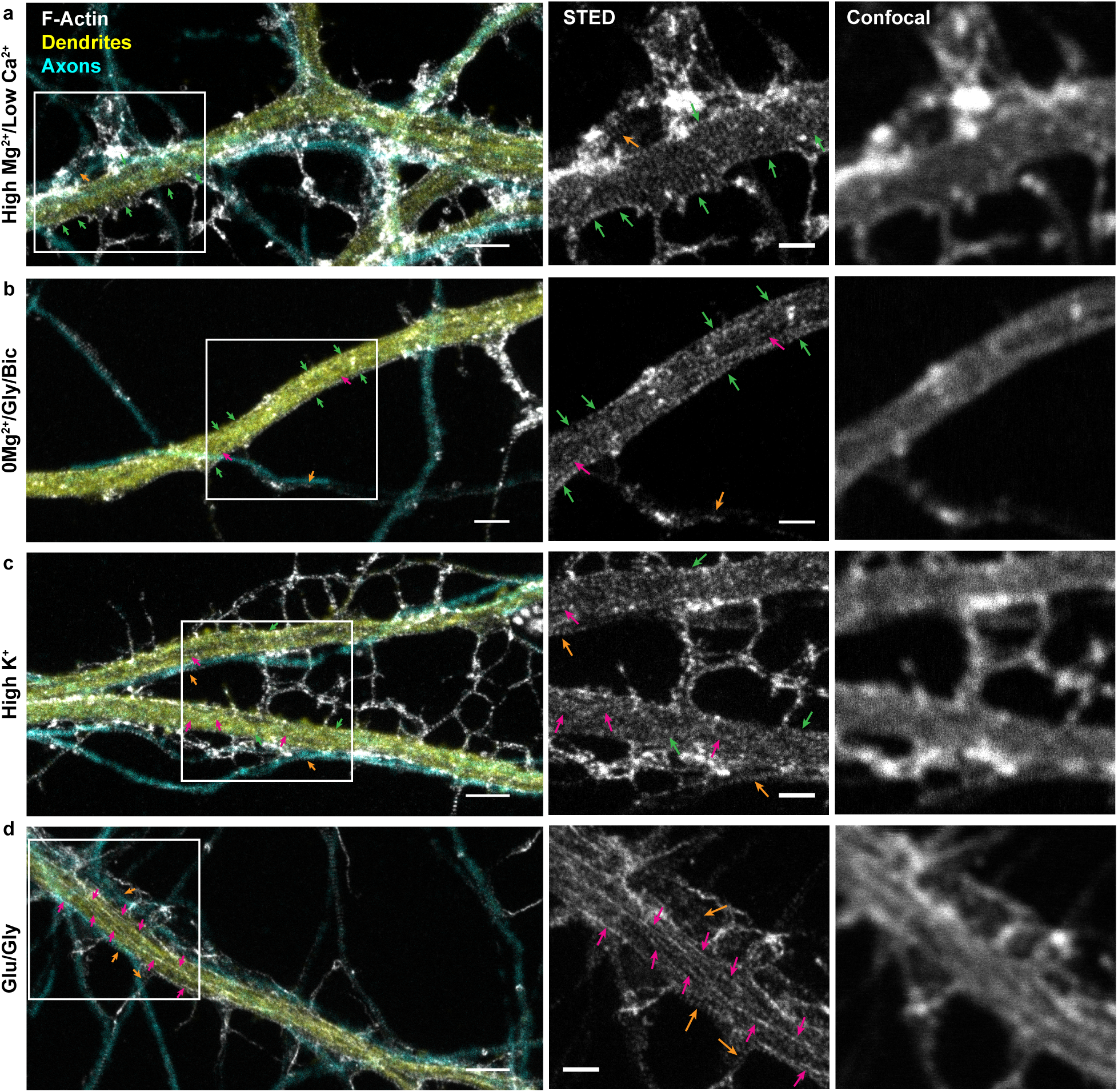
Nanoscale activity-dependent remodelling of F-actin revealed by STED nanoscopy. Three color imaging of F-actin (STED, white), phosphorylated neurofilaments (confocal, cyan) and MAP2 (confocal, yellow) was performed to identify F-actin nanostructures in dendrites (yellow) and axons (cyan). STED imaging shows that the prevalence of dendritic rings (green arrows) and longitudinal fibers (magenta arrows) is modulated by neuronal activity, while axonal rings (orange arrows) are observed regardless of the activity level. Shown are example images for a) the activity-reducing stimuli high Mg^2+^/low Ca^2+^ and the three activity-promoting stimuli b) 0Mg^2+^/glycine/bicuculline (0Mg^2+^/Gly/Bic), c) high K^+^, and d) glutamate/glycine (Glu/Gly). Insets (right) show a magnification of the regions identified with a white rectangle on the left STED images. Scale bar left: 2 *µ*m, insets: 1 *µ*m. For the raw images without overlay see Supplementary Fig. 3.

We next needed to apply a method to quantify the activity-dependent remodeling of the submembrane lattice. Several studies have examined the periodical pattern of *β*II-spectrin, which has been shown to be very clearly labelled in axons using *β*II-spectrin antibodies and readily quantifiable using autocorrelation routines [1, 3, 11]. In dendrites however, the concentration of *β*II-spectrin varies during developments [3, 4] and is more sparsely distributed in the dendritic lattice, compared to the axonal one [4, 12]. We nevertheless wanted to examine whether immunolabeling the neurons with *β*II-spectrin could help our quantification. We performed two-color STED nanoscopy of F-actin (Phalloidin-STAR635) and *β*II-spectrin (STAR580), in combination with confocal imaging of MAP2 (STAR488) to identify the dendrites. Hippocampal neurons (13 DIV), incubated in a high Mg^2+^/low Ca^2+^ to reduce neuronal activity, showed clear F-actin periodical ring patterns in dendrites, while the *β*II-spectrin pattern was less organized (Fig. 3a, green arrows). In contrast, the *β*II-spectrin periodical pattern in the axons was clearly detected (Fig. 3b, orange arrows).

**Figure 3:**
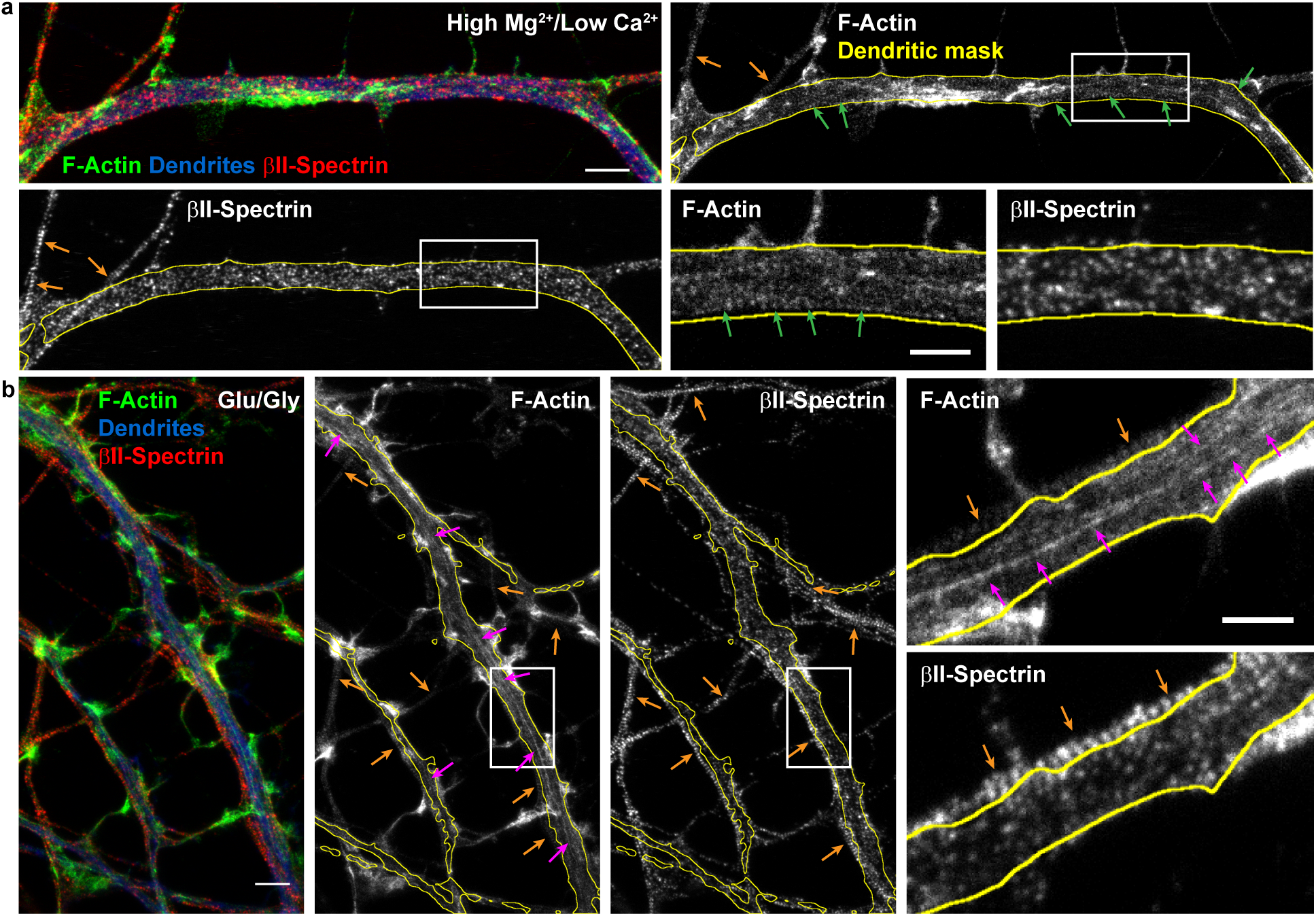
Two-color STED nanoscopy of F-actin and *β*II-spectrin in dendrites. (a) Low-activity high Mg^2+^/low Ca^2+^ condition and b) Glutamate/Glycine neuronal stimulation. *(a)Top-Left and (b)Left*: Overlay of two-color STED nanoscopy of F-actin (green) and *β*II-spectrin (red) with confocal imaging of MAP2 (blue) to identify the dendrites. Insets show the magnified regions identified in the full single-color STED images of F-actin and *β*II-spectrin. Arrows indicate the detected F-actin and *β*II-spectrin patterns: dendritic rings (green), axonal rings (orange), and longitudinal fibers (magenta). Scale bars (a, b) 2 *µ*m, insets 1 *µ*m.

We tested the strongest condition of activity-dependent remodelling of F-actin observed (Glu/Gly; Fig. 2d). While this treatment elicited a clear formation of longitudinal F-actin fibers in dendrites, the pattern of *β*II-spectrin appeared again disorganized. Meanwhile, in axons, the periodical *β*II-spectrin and F-actin patterns remained stable and clear after glutamate/glycine stimulation (Fig. 3b). Since the prevalence of the different spectrin isoforms (*β*II, *β*III, *β*IV) changes during development in dendrites and differs compared to axons[3, 4], it is possible that labelling only one spectrin subtype is not optimal to resolve the dendritic lattice organization. We thus elected to continue monitoring F-actin to quantify the activity-dependent remodeling of the submembrane lattice.

To characterize the prevalence of F-actin nanostructures in axons and dendrites, whole-image labelling has previously been applied [2]. We considered a quantitative approach, in which we manually labeled the F-actin rings or longitudinal fibers using polygonal bounding boxes to evaluate the remodelling of F-actin (Suppl. Fig. 4). However the diversity of patterns, highly variable across neurons, posed a significant challenge of time consuming manual annotation (average labeling time of 30 min per image). Furthermore, the large image sizes (1 × 10^6^ to 9 × 10^6^ pixels) made this process tedious and subject to decision fatigue [32], thereby limiting its application for testing several conditions. We thus decided to develop a high throughput analysis framework for the quantification of the activity-dependent F-actin reorganization in dendrites and axons.

**Figure 4:**
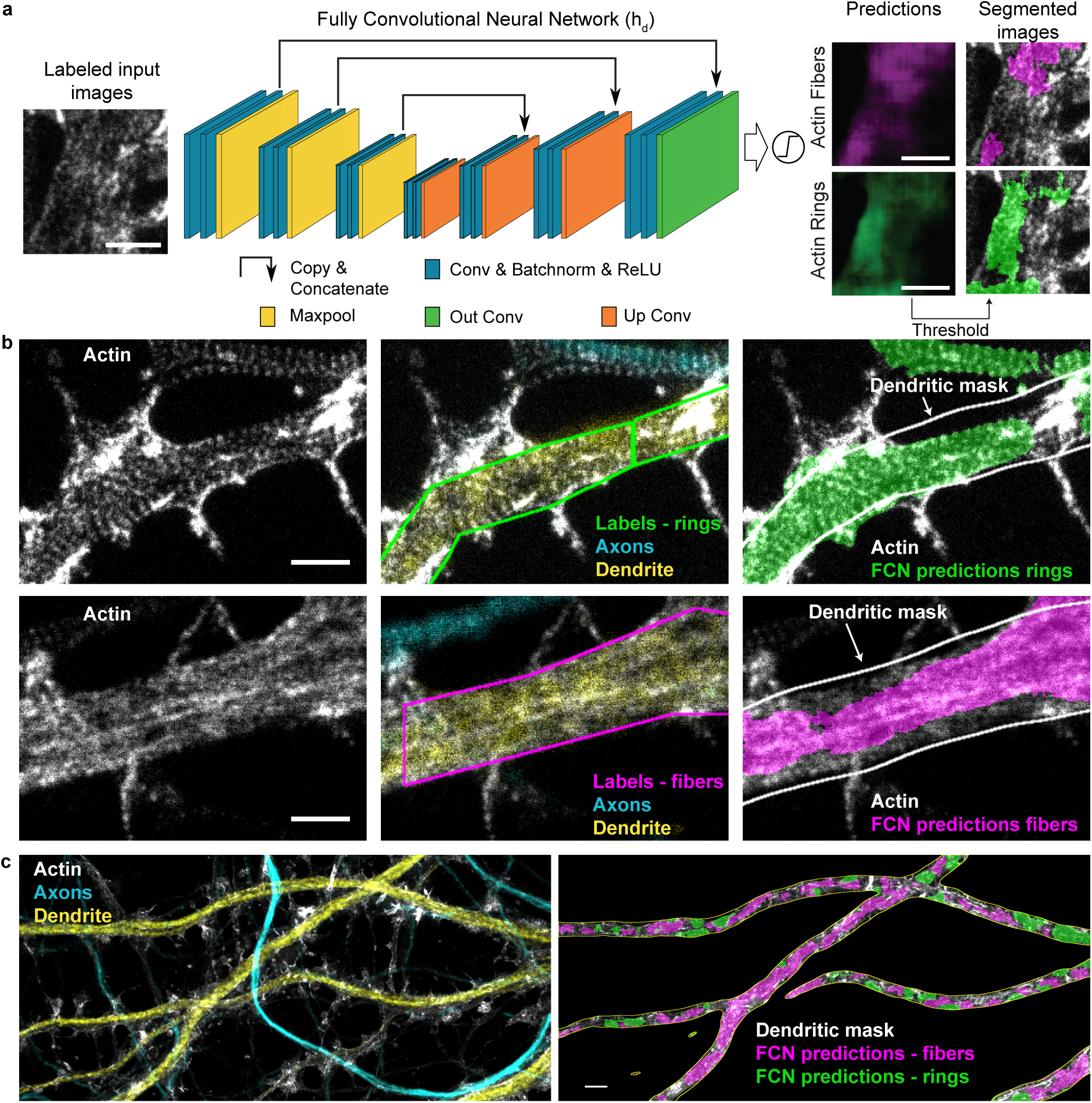
Segmentation of F-actin rings and longitudinal fibers using a fully convolutional neural network. a) Architecture of the fully convolutional network (FCN) (h_d_), which is a modified 2D U-Net (see Materials and Methods for specific implementation details, Conv: Convolution, Batchnorm: Batch Normalization, ReLU: Rectified Linear Unit). h_d_ is trained with images labeled for F-actin rings (green) and fibers (magenta). It generates scores between 0 and 1 for each pixel to create prediction maps for both structures. Independent thresholds are applied for rings (0.25) and fibers (0.4) to obtain two segmentation maps (see Materials and Methods and Supplementary Fig. 7). b) Comparison between the labeling of an expert (middle) and the corresponding FCN segmented image (right) on a representative image from the testing dataset. MAP2 (yellow) and phosphorylated neurofilaments (cyan) immunostaining and corresponding confocal images are used to identify dendrites and axons, respectively. Quantification of F-actin rings and fibers was performed within a dendritic mask generated from the MAP2 channel (white line, right). c) Representative input image analyzed with the FCN. The segmented area for F-actin rings (green) and fibers (magenta) is calculated inside the dendritic mask (white line) (right image) for each image. Scale bars (a, b) 1 *µ*m, (c) 2 *µ*m. For the raw images without overlay see Supplementary Fig. 5.

### Deep learning based analysis of F-actin nanostructures in axons and dendrites

To achieve reliable and high throughput quantification of the F-actin patterns at the nanoscale, we implemented a deep learning approach for the precise segmentation of F-actin rings and longitudinal fibers on STED images (Fig. 4). We chose to use a modified version of the U-Net architecture, a fully convolutional network (FCN), as it is known to perform well for biomedical image segmentation [15] (See Materials and Methods and Fig. 4a). Training such network generally requires a large amount of labeled data or the use of massive data augmentation [15]. However, the tediousness of the data labeling process of these complex F-actin patterns limited the amount of available data for FCN training. Meanwhile, data augmentation relies on the possibility to add new training samples by distorting or modifying existing samples in such a way that it does not alter their semantic interpretation. In the context of super-resolution microscopy, many of the usual alterations (stretching, noise addition, etc.) affect the spatial relation between fluorescent structures.

Considering these constraints, we formulated the segmentation task as a weak supervision problem to generate a sufficiently large dataset [20]. The expert had to label the F-actin patterns using polygonal bounding boxes instead of carefully drawing contours of each pattern (Fig. 4b, Supplementary Fig. 4), thereby reducing considerably the required labeling time to generate the necessary dataset. We trained a first FCN (see Materials and Methods) for the detection of F-actin rings and longitudinal fibers in dendrites (h_d_, Fig. 4, Supplementary Fig. 5). A second one, h_a_, was trained to detect solely F-actin rings in axons since we did not observe longitudinal fibers in axons regardless of the experimental condition (h_a_, Supplementary Fig. 6a). We characterized the performance of both FCNs using standard metrics such as F1-score, specificity, sensitivity and precision (see subsection on Performance Metrics in the Material and Methods, Supplementary Fig. 6, 7). To assess the suitability of this method for the quantitative analysis of the ratios between F-actin rings and longitudinal fibers in dendrites (Fig. 4c), we compared detected areas obtained by expert or h_d_ labeling (Supplementary Fig. 7c). The results show no significant difference in the F-actin rings and fibers ratio detected by h_d_ and an expert (Supplementary Fig. 7c).

Fig. 5a,b demonstrate that h_d_ can use these labels to learn identifying the structures of interest, while generating more accurate segmentation compared to the coarse labels it was provided with. The precision-recall curves (Fig. 5b) were obtained from the comparison of the FCN predictions and bounding box annotations with a precisely annotated dataset, which is considered as the ground truth (P, see Materials and Methods). We observe a higher precision for h_d_ segmentation over all range of recall compared to polygonal bounding box annotations, implying that the labels generated by the network are closer to the ground truth than the manual bounding box annotations. Without learning precise segmentation rules, the system performance would be upper-bounded by the quality of the segmentation provided by a polygonal bounding box on P. This ability to cope with less precise and missing information is a key element in the application of deep learning methods for high throughput bio-imaging segmentation tasks, where the acquisition of more or better labeled images can be impracticable. To better characterize this ability, we designed two experiments. We first tested the sensitivity of h_d_ to omission of ground truth labels in the training dataset (polygonal bounding boxes), since we observed that manual labeling of large microscopy images was prone to this type of labeling errors. To this extent, we removed up to 70% of the expert labels in the training dataset, while keeping the number of training crops constant (See Materials and Methods and Supplementary Fig. 9). We observed that the performance of h_d_ is only slightly affected by sparse expert labeling and that even removing one out of two labels on the F-actin patterns resulted in a precision comparable to the original one (Supplementary Fig. 9a,b).

**Figure 5:**
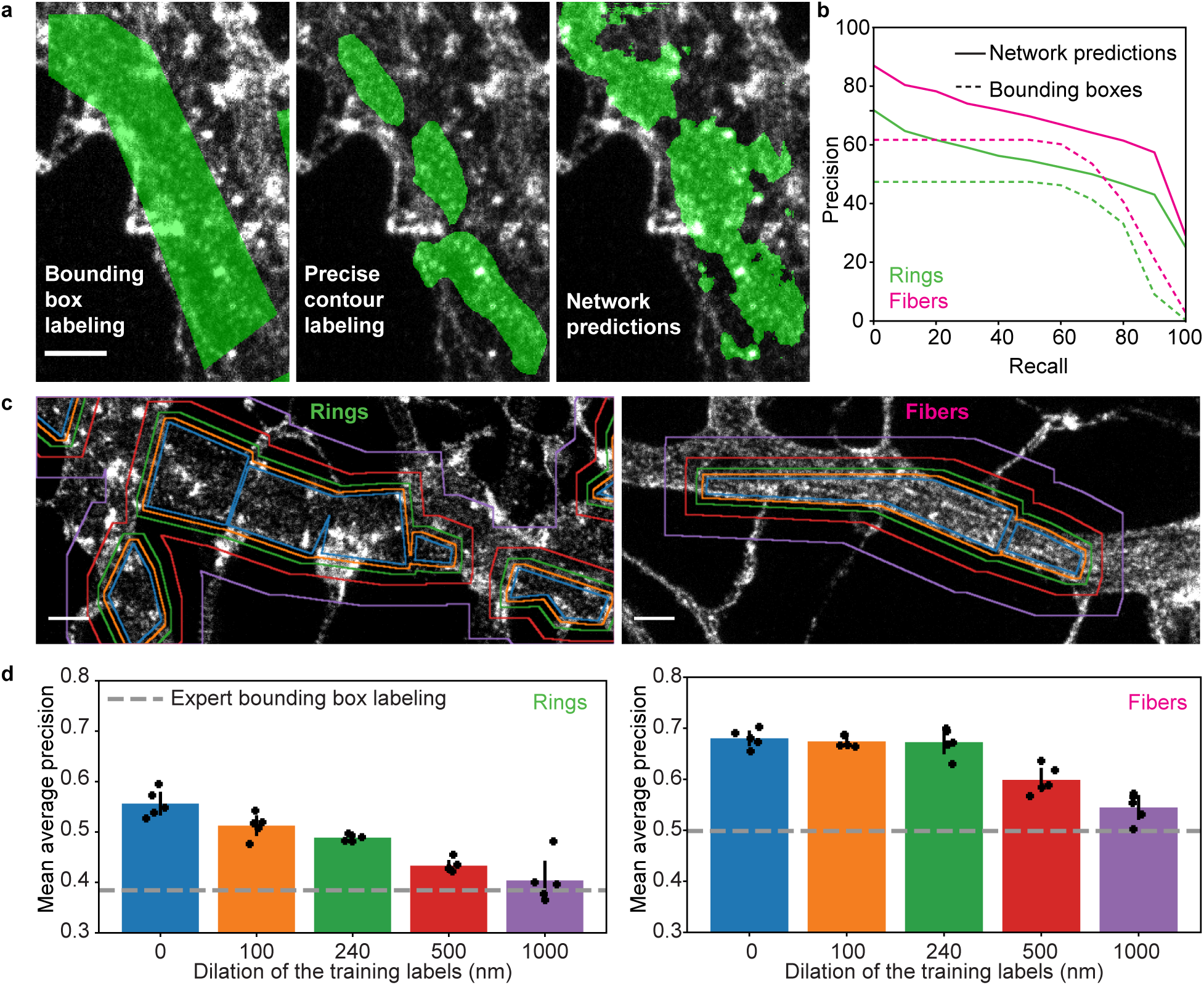
Performance evaluation of the FCN h_d_ in the context of weakly supervised learning and labeling errors. a) Comparison between manual bounding box (left), precise contour (middle) and FCN predicted (right) labeling of the dendritic F-actin ring pattern (For the raw image without overlay see Supplementary Fig. 8). b) Precision-recall curves for F-actin rings (green) and longitudinal fibers (magenta). The area under the curve, or average precision (AP), was calculated for both patterns. The network achieved a AP score of 0.53 and 0.67 for F-actin rings and fibers respectively compared to 0.38 and 0.5 for the manual bounding box labeling (using the precise contour labeling as the ground truth). The higher performance observed for the predictions compared to the bounding box labeling shows that the network is able to infer precise segmentation rules using only coarse examples. c) Generation of a training dataset to characterize the impact of coarse labeling on the precision of h_d_ by stepwise dilation (original labels - blue, 100 nm - orange, 240 nm - green, 500 nm - red, 1 *µ*m - violet) of the training labels for F-actin rings (left) and fibers (right). d) The AP scores were calculated for 5 different instances of the network for each dilation step. For F-actin fibers (right) dilation up to 1 *µ*m still resulted in network predictions with significantly higher precision than manual bounding box labeling (post-hoc t-test, *p*_original_ = 1.8210 × 10^−6^, *p*_100nm_ = 2.4291 × 10^−7^, *p*_240nm_ = 1.8530 × 10^−5^, *p*_500nm_ = 2.7931 × 10^−4^, *p*_1*µ*m_ = 9.7314 × 10^−3^). For the F-actin ring patterns, a dilation of 1 *µ*m led to comparable AP than the expert labeling (post-hoc t-test, *p*_1*µ*m_ = 0.3309), while smaller dilation steps led to significantly higher AP scores compared to bounding box labeling (post-hoc t-test, *p*_original_ = 1.9383 × 10^−5^, *p*_100nm_ = 4.1884 × 10^−5^, *p*_240nm_ = 2.2539 × 10^−7^, *p*_500nm_ = 2.3219 × 10^−4^). Black lines represent the 95% confidence interval calculated from the t-statistics distribution. Scale bars 1 *µ*m.

We also characterized how coarse labeling would influence the precision of the network. To this extent, we carefully labeled a small dataset of 70 images to calculate the average precision score (AP) of manual bounding box labeling (Fig. 5d, gray dotted line). We next compared the AP of manual bounding box labeling to the performance of h_d_ trained with increasing label size (obtained by stepwise uniform dilation of the bounding box labels, see Materials and Methods, Fig. 5c,d). Our results show that even 1 *µ*m dilation of the training labels for the F-actin fibers and 500 nm dilation for the F-actin rings still lead to a significantly higher AP score compared to manual bounding box labeling (Fig. 5d). We thus concluded that our weakly supervised deep learning approach was reliable to analyse quantitatively the remodeling of F-actin patterns at the nanoscale in neurons.

### Quantification of the activity-dependent reorganization of the F- actin lattice in dendrites and axons using deep learning

We first used the FCN h_a_, trained on bounding boxes identifying axonal F-actin rings, to quantify the presence of this structure in axons for different levels of neuronal activity (Fig. 6a,b, Supplementary Fig. 10). Using this approach, we measured the area of detected F-actin rings within axons, co-stained with SMI31 to generate an axonal mask (mean total length per image of 151*±*89*µ*m and maximal branch length of 32 *±* 13*µ*m, Fig. 6a, Supplementary Fig. 11a,b). This allowed to automatically quantify the proportion of F-actin rings in axons on a large dataset and for different neuronal activity levels (150 neurons, 4 independent cultures, 4 experimental conditions). Our quantitative approach reliably detected an F-actin periodical lattice in all analysed images of axons and revealed no activity-dependent change in its prevalence (one-sided ANOVA, p = 0.2525, Fig. 6b).

**Figure 6:**
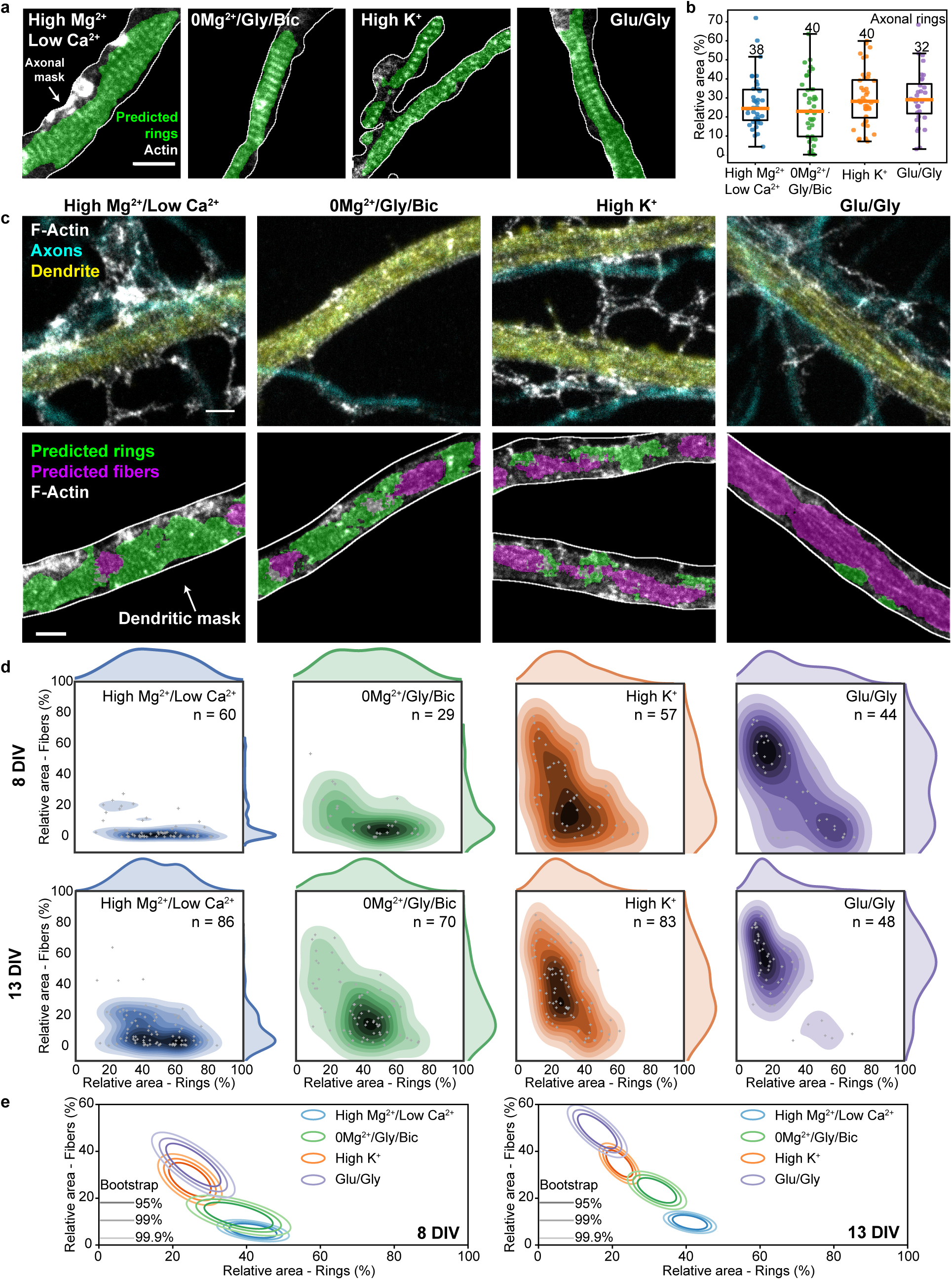
Increasing neuronal activity induces the reorganization of F-actin rings into longitudinal fibers in dendrites but not in axons. a) Representative images of the periodical F-actin rings in axons of neurons exposed to 4 different treatments modulating neuronal activity. The area of F-actin rings segmented by the FCN h_a_ is shown in green. b) The detected area for axonal rings in 13 DIV neurons remains unchanged upon stimulation (one-sided ANOVA, *p* = 0.2525, numbers above each boxes indicate the number of neurons from 4 independent cultures). c) Representative images of the periodical F-actin lattice and the longitudinal fibers in dendrites of 13 DIV neurons for four different treatments modulating neuronal activity: *Top* STED Images of F-actin stained with Phalloidin-STAR635 overlayed with the confocal images of the dendritic (MAP2, yellow) and axonal (phosphorylated neurofilaments, cyan) makers. *Bottom*: Predictions of the FCN for F-actin rings (green) and fibers (magenta) inside the dendritic mask (white line). d) Bivariate kernel density estimate of the raw data (grey cross) for 8 DIV and 13 DIV neurons treated with i) high Mg^2+^/low Ca^2+^ for 10 min (blue), ii) 0Mg^2+^/Gly/Bic for 10 min (green), iii) high K^+^ for 2 min (orange) and iv) Glu/Gly for 2 min (violet). e) Mean distributions using bootstrapping. The formation of F-actin fibers is enhanced for 13 DIV neurons compared to 8 DIV neurons for the synaptic stimulation 0Mg^2+^/Gly/Bic or Glu/Gly, but not for high K^+^ stimulations. Shown are the regions comprising 95%, 99% and 99.9% of the data point distribution. Scale bars 1 *µ*m. d,e) Number of independent cultures (N): high Mg^2+^/low Ca^2+^ *N*_8DIV_ = 6, *N*_13DIV_ = 9; 0Mg^2+^/Gly/Bic *N*_8DIV_ = 4, *N*_13DIV_ = 8; high K^+^ *N*_8DIV_ = 6, *N*_13DIV_ = 9; Glu/Gly *N*_8DIV_ = 5, *N*_13DIV_ = 6. Note that the detected areas for dendrite and axons cannot be compared since the detection of F-actin rings was performed with two different FCNs and using different foreground masks. Only a comparison between the stimulation conditions for each experiment (dendrites or axons) is possible. Scale bars 1 *µ*m. For the raw image without overlay see Supplementary Fig. 10

By contrast, our deep-learning based approach revealed significant effects on the F-actin patterns in dendrites of 13 DIV neurons (we also observed a clear activity-dependent remodeling in the soma, but did not quantify it, Supplementary Fig. 1). Automated analysis of the F-actin patterns within a dendritic mask was performed using the MAP2 confocal signal (mean total length per image of 71 *±* 36*µ*m and maximal branch length of 42 *±* 10*µ*m, Fig. 6c, Supplementary Fig. 11c,d). The analysis revealed a remodelling of the F-actin rings into longitudinal fibers, scaling with the strength of the activity promoting stimuli (Fig. 6c). Under reduced neuronal activity (high Mg^2+^/low Ca^2+^), the dendritic F-actin rings were most prevalent, while longitudinal fibers were rarely detected (Fig. 6c-e). We used a high Mg^2+^/low Ca^2+^ solution to reduce neuronal activity as the variability between neurons in the prevalence of F-actin rings and fibers was reduced compared to untreated cells (see Supplementary Fig. 12). Promoting synaptic activity with 0Mg^2+^/Gly/Bic significantly reduced the proportion of dendritic area exhibiting F-actin rings and increased the prevalence of longitudinal fibers (*p* = 2.9 × 10^−6^, randomization test (RdT, see Materials and Methods)). The ring pattern was even more strongly reduced with brief (2 min) high K^+^ or Glu/Gly stimulation ((RdT: *p*_highK_+ = 9.1 × 10^−15^, *p*_Glu*/*Gly_ = 0, Fig. 6c-e). Meanwhile, we observed a correlative progression of the formation of F-actin longitudinal fibers with the strength of neuronal activity stimulation (Fig. 6c-e). These F-actin fibers did not colocalize with microtubules (Supplementary Fig. 13), suggesting that both structures are not directly associated. These results indicate that while neuronal activity has no impact on the stability of the F-actin submembrane lattice in axons, it causes its reorganization into longitudinal fibers in dendrites.

Since the F-actin/spectrin lattice has been reported to be regulated by neuronal development in dendrites [2, 4], we compared the effects of the different stimuli on 8 and 13 DIV neurons (Fig. 6d,e). The results indicated that dendritic F-actin rings were slightly more prevalent, whereas fibers were less prevalent, in young neuron compared to older ones, at low neuronal activity (high Mg^2+^/low Ca2+). Synaptic stimulation (0Mg^2+^/Gly/Bic) elicited a significantly stronger F-actin dendritic reorganization in mature cultures, whereas broad membrane depolarization with high K^+^ caused a similar extent of reorganization at both ages (RdT: 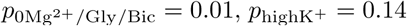). Finally, Glu/Gly stimulation caused a strong remodeling of F-actin in young neurons and a nearly complete transition to dendritic fibers in mature cultures (RdT: *p*_Glu*/*Gly_ = 1.1 × 10^−4^, Fig. 6d,e). These results indicate that the activity-dependent remodeling of dendritic F-actin cytoskeleton is more pronounced in mature and more synaptically connected neurons.

To test whether the impact of the 0Mg^2+^/Gly/Bic stimulation depended on action potential firing, we tested the effect of co-application of the sodium channel blocker tetrodotoxin (TTX). We observed a partial but significant reduction in F-actin remodeling by TTX (RdT: *p* = 1.4 × 10^−3^, Fig. 7a,b). It was shown that miniature excitatory potentials can drive NMDA receptor (NMDAR) activity in absence of Mg^2+^ [33]. We thus tested the effect of NMDAR blocker (2R)-amino-5-phosphonovaleric acid (APV) on the 0Mg^2+^/Gly/Bic stimulus and found that it reduced more strongly the F-actin remodeling (RdT: *p* = 2.5 × 10^−5^, Fig. 7a-c, Supplementary Fig. 14), compared to TTX. These results indicate that synaptic NMDAR activity can trigger F-actin remodeling from a ring to fiber pattern in dendrites.

**Figure 7:**
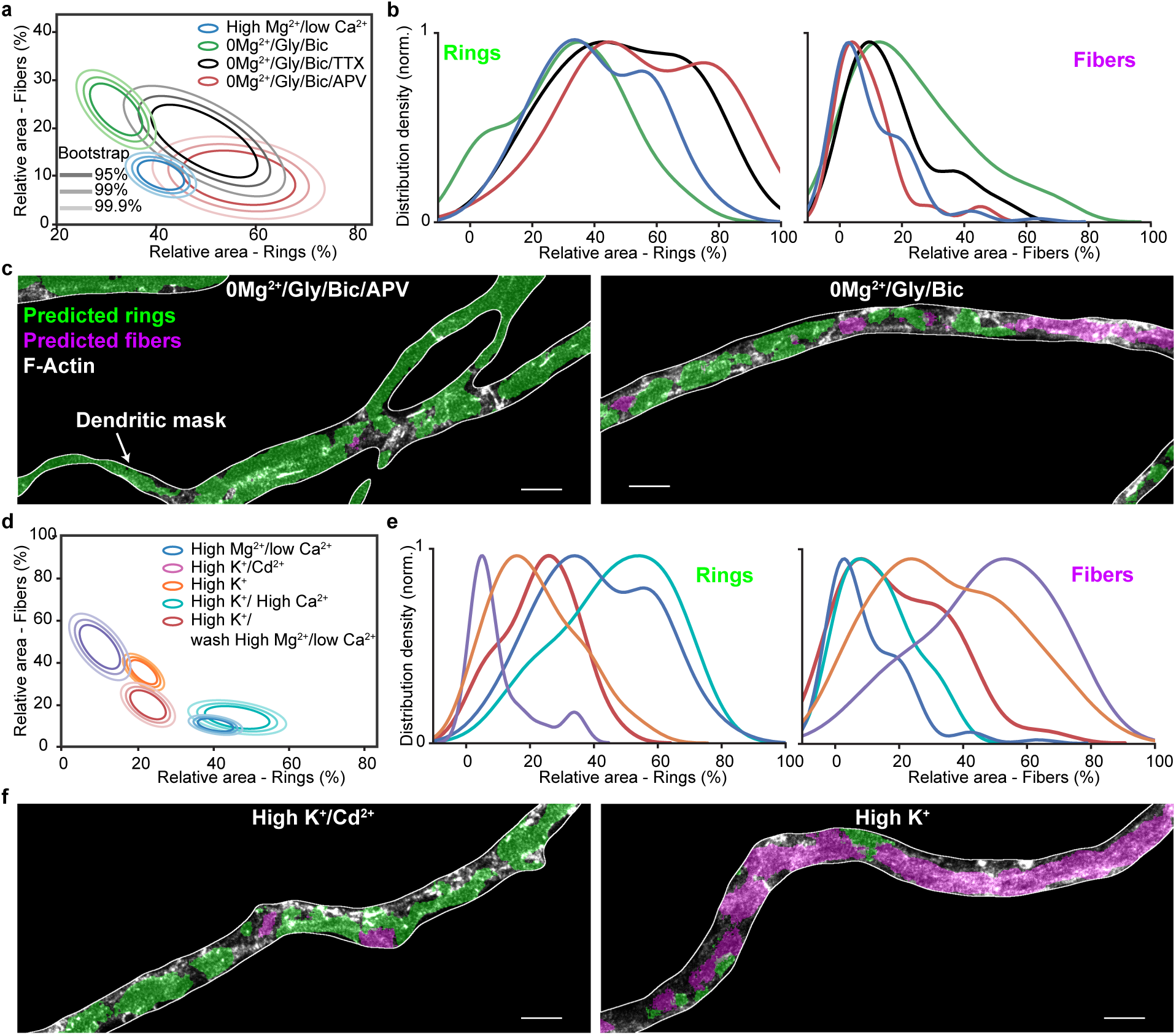
Synaptic NMDAR activity and Ca^2+^ influx can drive a reversible dendritic F-actin reorganization from a ring to fiber pattern. a) Mean distributions of dendritic F-actin rings and fibers using bootstrapping for synaptic stimulation (0Mg^2+^/Gly/Bic for 10 min) without (green) or with TTX (violet) or APV (red) compared with the low activity high Mg^2+^/low Ca^2+^ condition (blue). b) Density distribution of the raw data. TTX (1 *µ*M) partially but significantly blocks the F-actin remodeling caused by 0Mg^2+^/Gly/Bic stimulation, while APV (25 *µ*M) blocks it further (*p* = 1.4 × 10^−3^ and *p* = 2.5 × 10^−5^ respectively). c) Representative images of neurons treated with 0Mg^2+^/Gly/Bic with (left) and without (right) APV segmented with our deep learning based approach. d) Mean distributions of dendritic F-actin rings and fibers using bootstrapping for 2 min high K^+^ stimulation (1.2 mM Ca^2+^, orange) or with 2.4 mM Ca^2+^ (violet) or with 1.2 mM Ca^2+^ and 50 *µ*M Cd^2+^ (pink). The red circles indicate the same high K^+^ stimulation (1.2 mM Ca^2+^) condition followed by 15 min wash in high Mg^2+^/low Ca^2+^. The F-actin remodeling is Ca^2+^-dependent and reversible, at least partially within 15 min. e) Density distribution of the raw data. f) Representative images of neurons treated with high K^+^ stimuli with (left) and without (right) Cd^2+^ segmented with our deep learning based approach. Statistical analysis performed with a randomization test (see Materials and Methods). Number of independent cultures (N) and number of neurons (n): high Mg^2+^/low Ca^2^+ *N* = 9, *n* = 86, 0Mg^2+^/Gly/Bic *N* = 8, *n* = 70; 0Mg^2+^/Gly/Bic 1*µ*M TTX *N* = 2, *n* = 20; 0Mg^2+^/Gly/Bic + 25*µ*M APV *N* = 2, *n* = 20; high Mg^2+^/low Ca^2+^ *N* = 9; high K^+^/1.2 mM Ca^2+^ *N* = 9, *n* = 83; high K^+^/2.4 mM Ca^2+^ *N* = 2, *n* = 20; high K^+^/50 *µ*M Cd^2+^ *N* = 2, *n* = 22; high K^+^/15 min wash high Mg^2+^ *N* = 3, *n* = 32. For the raw images without overlay see Supplementary Fig. 14

Neuronal depolarization and NMDAR activities drive Ca^2+^ influx, which activates a wide range of dendritic signaling processes. We thus assessed whether Ca^2+^ influx was mediating the high K^+^-induced remodeling of dendritic F-actin cytoskeleton, by either i) increasing the Ca^2+^ concentration or ii) blocking Ca^2+^ entry with Cd^2+^. In comparison to regular Ca^2+^ concentration (1.2 mM), 2.4 mM caused a significantly stronger reduction in F-actin rings accompanied by an increase in F-actin fibers (RdT: *p*_2.4*mMCa*_2+ = 6.6 × 10^−5^, Fig. 7d-f). Application of Cd^2+^ during the stimulation prevented the activity-dependent reorganization of the F-actin cytoskeleton; no significant reorganization of the F-actin cytoskeleton was observed compared to high Mg^2+^/low Ca^2+^ condition (RdT: *p*_*Cd*_2+ = 0.06, Fig. 7d-f, Supplementary Fig. 14). These results indicate a clear Ca^2+^-dependence for the activity-dependent F-actin remodeling in dendrites.

Finally, we assessed whether the F-actin remodelling was reversible by washing the neurons with high Mg^2+^/low Ca^2+^ activity blocking solution for 15 min after a 2 min high K^+^ stimulation. We observed a partial recovery of the F-actin rings combined with a strong reduction in F-actin fibers following the 15 min wash. These results suggest that the activity-dependent remodeling of F-actin is at least partially reversible (*p* = 3.1 × 10^−4^, Fig. 7d,e).

## Discussion

Our results demonstrate that the somatodendritic F-actin-based lattice is highly dynamic compared to that of the axon. While it was known that the dendritic lattice is distributed in a more patchy fashion, we show here that this difference is likely due, at least partially, to a distinct response to neuronal activity, which destabilizes the lattice only in dendrites and somata (Fig. 8). The sporadic presence of the submembrane lattice previously reported in dendrites may thus be due to variable levels of electrical or synaptic activities in the neuronal culture preparations, which have not been specifically controlled and which also depend on the developmental stage of the preparation [1, 2, 4, 12]. In our study, we specifically controlled both the neuronal activity level and the developmental stage of the preparation and found that the extent of dendritic remodeling is more pronounced in more mature and synaptically connected neurons. Another source of variability may have arisen from inconsistent use of specific axonal and dendritic markers to discriminate these processes. We found it critical to have double labeling of both axonal and dendritic markers in our cultures, as these distinct processes exhibit considerable spatial overlap, while their F-actin-based membrane lattice reacts differently to neuronal stimulation.

**Figure 8:**
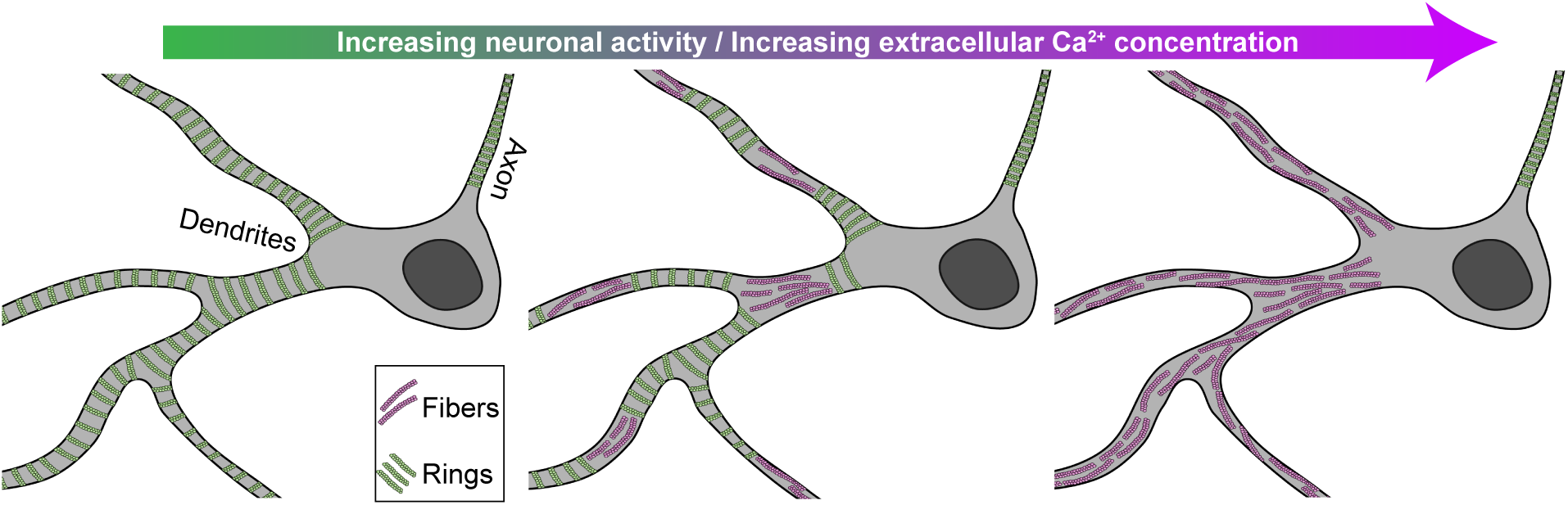
Schematic representation of the nanoscale patterns of F-actin cytoskeleton in axon and dendrites in a neuron undergoing variable level of neuronal activity; the periodical lattice reorganizes into longitudinal fibers in dendrites, but not in axons, with increasing neuronal/Ca^2+^ activity.

We demonstrated that the activity-dependent disappearance of the somatodendritic submembrane lattice coincided with the formation of F-actin longitudinal fibers. The simplest interpretation of this observation is that the F-actin underwent restructuring from a ring to fiber pattern. This putative remodeling would be demonstrated more clearly using live imaging, however, as we described and discussed above, currently available F-actin probes either do not reveal, or interfere with, the F-actin lattice dynamics. Nevertheless, the Ca^2+^-dependence of this proposed remodeling is consistent with previous work showing that nucleation, capping, and severance of F-actin is regulated by Ca^2+^ [34]. Meanwhile the lattice remodeling might also be modulated by spectrin cleavage through Ca^2+^-dependent calpain [11, 35]. The reasons why the axonal lattice is resistant to enhanced neuronal activity remain to be determined. The elevation in axonal Ca^2+^ may be reduced compared to other domains. Alternatively, the composition of the lattice may differ in axons, making it less sensitive to Ca^2+^-mediated destabilisation. The selective presence in axons of transmembrane and extracellular matrix-interacting proteins, such as neurofascin, which has been shown to intercalate with F-actin rings in axons [2], may have a stabilizing role for the submembrane lattice.

The stability of the membrane-associated cytoskeleton was shown to be important for RTK signaling [11]. The activity-dependent remodeling of the F-actin based lattice into F-actin longitudinal fibers might, on the other hand, be necessary for some dendritic signaling mechanisms, such as endo/exocytosis, membrane receptor lateral movement, intracellular trafficking or spine formation. In contrast, the stable lattice in axons might limit transmembrane receptor movement [36], which could support a tight location of voltage-dependent channels critical for action potential conduction. It might also serve to restrict exo/endocytosis to pre-synaptic terminals, where the lattice is less organized [2, 12]. In contrast, the removal of the lattice in dendritic segments may release a break on membrane receptor diffusion and exo/endocytosis, or spine formation. Meanwhile, the activity-dependent formation of F-actin longitudinal fibers in dendrites may serve to promote active cargo transport [5, 37] and targeting of material to synapses [7]. As such, the activity-dependent conversion of membrane-associated rings to intracellular fibers of the dendritic F-actin cytoskeleton may be essential for synaptic plasticity. In this context, we found that NMDA receptor function, which is crucial for synaptic plasticity, was essential in supporting the remodeling of the dendritic F-actin rings to longitudinal fibers. Hence, of the numerous signaling processes downstream of NMDA receptor function and Ca^2+^ during synaptic plasticity, the remodeling of the dendritic F-actin-based lattice into longitudinal fibers may be of significance.

The quantification of these fluorescent patterns is highly difficult because of the heterogeneity and complexity of the signals, and because it is prone to biases. Our study emphasizes the potential of machine learning approach, as an expert surrogate, for high throughput analysis of complex and highly variable neuronal structures observed with optical nanoscopy. It became essential in order to pick up subtle differences associated with developmental stages, activity levels or drug action. The segmentation task shown here presented a number of challenges related to the context of experimental biology: 1) limited number of biological samples and labeled data (keeping in mind that increasing the amount of images for the training dataset reduces the available dataset for biological analysis), 2) high variability across samples, 3) expert-dependent determination of the ground truth, 4) no direct link to other problems that could be used as a pre-training step. We demonstrated how a deep learning approach can be applied, using weak supervision and harnessing the deep network extrapolation power. We also characterized how two types of labeling problems (e.g. incomplete and decreased precision) influence the performance of the FCN in this weakly supervised learning framework. We showed that our approach was sufficiently robust to the use of coarse labeling and label omission. This should broaden the possibilities of exploiting deep learning-based analysis for high throughput biomedical image analysis, especially when the generation of a large precisely labeled training dataset is impracticable. The weakly supervised deep learning approach used in our study provided the necessary tool to investigate more effectively and accurately the activity-dependent remodeling of the dendritic F-actin cytoskeleton, and should be useful for similar types of optical imaging-based investigations of cellular signaling.

## Materials and Methods

### Cell culture and neuronal stimulations

Neuronal cultures were prepared from neonatal Sprague Dawley rats. We followed the guidelines of the animal care committee of Université Laval for the care and use of the rats. Before dissection of hippocampi, animals were sacrificed by decapitation, in accordance to the procedures approved by the animal care committee of Université Laval. Dissociated cells were plated on poly-d-lysine coated glass coverslips (∼ 12mm) at a low density of 25 000 cells/ml. The growth medium consisted of Neurobasal and B27 (50:1), supplemented with penicillin/streptomycin (50 U per mL; 50 *µ*g per mL) and 0.5 mM L-GlutaMAX (Invitrogen). High density neurons (10 million cells/mL) were plated directly in poly-d-lysine coated wells to serve as a feeding layer for the low density coverslip, placed upside down above the feeding layer. To limit proliferation of non-neuronal cells, Ara-C (2.5 *µ*M,; Sigma-Aldrich) was added to the media 2 days later. Thereon, the culture was fed twice a week by replacing ∼1/3 of the growth medium with serum- and Ara-C–free medium.

Neuronal stimulations were performed in HEPES buffered solutions at 37°C. The following solutions were used: high Mg^2+^/low Ca^2+^ (in mM: NaCl 98, KCl 5, HEPES 10, CaCl_2_ 0.6, Glucose 10, MgCl_2_ 5), 0Mg^2+^/Gly/Bic (in mM: NaCl 104, KCl 5, HEPES 10, CaCl_2_ 1.2, Glucose 10, MgCl_2_ 0, Glycine 0.2, Bicuculline 0.01), high K^+^ (in mM: NaCl 75, KCl 40, HEPES 10, CaCl_2_ 1.2, Glucose 7.5, MgCl_2_ 1), Glu/Gly (in mM: NaCl 102, KCl 5, HEPES 10, CaCl_2_ 1.2, Glucose 10, MgCl_2_ 1; Glutamate 0.1, Glycine 0.01); Osmolality: 240-250 mOsm/kg, pH: 7.35. Incubation lasted 10 min for high Mg^2+^ and 0Mg^2+^/Gly/Bic treatments and 2 min for high K^+^ and Glu/Gly stimulation. After the treatment, the cells were directly transferred in a 4% paraformaldehyde (PFA) solution for fixation (See Fixation and Immunostaining). To characterize the impact of action potential firing or NMDAR activity on F-actin reorganization during the synaptic stimulation 0Mg^2+^/Gly/Bic, 1*µ*M TTX or 25 *µ* APV were added, respectively, to the stimulation solution. To assess the reversibility of the actin reorganization caused by high K^+^ stimulation, cells were first stimulated 2 min in high K^+^ solution and transferred into 37°high Mg^2+^/low Ca^2+^ blocking solution for 15 min prior to fixation. The effect of Ca^2+^ on the F-actin reorganization was assessed either by increasing the Ca^2+^ concentration of the high K^+^ solution (CaCl_2_ 2.4 mM) or by blocking calcium channels with Cd^2+^ (100*µ*M CdCl_2_) during high K^+^ stimulation.

Latrunculin A was used to disrupt the F-actin lattice as reported previously [4]. Latrunculin A was added to the culture media for 1 h with a final concentration of 10*µ*M. Subsequently, cells were incubated 10 min in high Mg^2+^/low Ca^2+^ solution with 10 *µ*M latrunculin A and fixed with 4% PFA solution.

### Fixation and Immunostaining

Cultured hippocampal neurons were fixed in 4% PFA solution, permeabilized with 0.1% Triton X-100, blocked with 2% goat serum and immunostained as described previously [38]. To improve the F-actin staining, phalloidin was incubated for 2h following the immunostaining steps with primary and secondary antibodies. Coverslips were mounted in Mowiol-DABCO for imaging. F-actin was stained with phalloidin- STAR635 (Abberior, cat. 2-0205-002-5, 1:50 dilution). Dendrites were identified using a staining against the microtubule-associated-protein-2 with Rabbit-anti-MAP2 PAB (Milipore Sigma, cat. AB5622, 1:1000) and GAR-STAR488 SAB (Abberior, cat. 2-0012-006-5, 1:250). Axons were identified using a staining against the phosphorylated neurofilaments (SMI31) stained with the PAB mouse-anti-SMI31 (Biolegend, cat. 801601, 1:250) and the SAB GAM-STAR580 (Abberior, cat. 2-0002-005-1, 1:250).

### STED-imaging

Super-resolution imaging was executed on a 4 color Abberior Expert-Line STED system (Abberior Instruments GmbH, Germany). Imaging of F-actin stained with Phalloidin-STAR635 was performed using a 640 nm pulsed diode (40 MHz), a 775 nm depletion laser (40 MHz) and a a ET685/70 (Chroma, USA) fluorescence filter. MAP2-STAR488 and SMI31-STAR580 were imaged in confocal mode using excitation diodes at 485 nm and 561 nm (40MHz), 525/50 and 615/20 (Semrock, USA) fluorescence filters respectively. Scanning was conducted in a line scan mode with a pixel dwell time of 15*µ*s and pixel size of 20 nm. For the STED image of the F-actin cytoskeleton a line repetition of 5 was selected. The microscope was equipped with a 100x 1.4NA, oil objective and fluorescence was detected on independent avalanche photodiode detectors (APD) with approximately 1 Airy unit detection pinhole. Our STED microscope was equipped with a motorized stage and auto-focus unit. Images were processed using FIJI (ImageJ) software [39].

### Dataset

For the training of the FCNs presented in this work, two labeled datasets (1. Dendritic F-actin rings and fibers and 2. Axonal rings) were built using a custom labeling application developed in Python.

For the segmentation of dendritic F-actin ring and fiber regions, we collected a set of large STED images (between 500 × 500 and 3000 × 3000 pixels) of the F-actin cytoskeleton. Each image contained 3 channels: 1) F-actin (phalloidin-STAR635), 2) an axonal marker (SMI31-STAR580), and 3) a dendritic marker (MAP2- STAR488). Areas exhibiting F-actin rings and fibers were labeled by an expert on the MAP2-positive regions (Fig. 4b). The labels were additionally filtered by keeping only the regions belonging to the foreground mask generated with the MAP2 channel (Otsu thresholding operation [40] applied after a gaussian blur with σ = 20 pixels). The training, validation, and testing datasets for dendritic F-actin structures consisted in respectively 42, 52, and 105 representative images, with neurons at different activity levels in all sets. The large testing dataset was designed to include a representative distribution of images from 8 different treatment conditions as well as sufficient number (12-15) of independent cells for each condition. We could therefore compare the performance of the FCN to the manual labelling for various activity levels, ensuring that no bias was induced by FCN labelling between different treatments.

While a FCN could be trained on large images, it is more convenient to use smaller crops for this purpose, since the model is size independent. We thus used a 128 × 128 sliding window with a 16 pixels overlap on the input images to generate a total of 2263 crops for the training set. To reduce class imbalance, we considered only crops comprising a label (rings or fibers) on at least 1% of their total area. The intensity in each crop was scaled in the [0, 1] range. For the scaling, the minimum and maximum values of each image in the dataset were computed. The median of all minima (`m`) was used as [0] and the median of all maxima plus 3 standard deviation (`M`) was used as [1]. This [`m, M`] interval was multiplied by a factor of 0.8 for the normalization. The resultant crop was clipped in the [0, 1] range to ensure the proper intensity scale for any input. Data augmentation was applied at training time (flip up-down and left-right, intensity scale and gamma adaptation), with a 50% probability for each operation.

For the segmentation of the periodical F-actin lattice in axons, a training set was built with 335 small images (224 × 224 pixels) of axons only stained with phalloidin-STAR635. The validation and testing set consisted in 112 and 69 images respectively. The axonal F-actin rings were labeled by an expert as described above. Sliding window, scaling and data augmentation operations were applied similarly to the dendritic dataset. For this dataset, we did not enforce any minimum labeled area.

### Fully Convolutional Neural Network

Two FCNs were implemented (h_d_, h_a_) based on the well-known U-Net architecture (Fig. 4a and Supplementary Fig. 6a) [15] using the PyTorch library.

The first FCN (h_d_) was designed for the detection of dendritic F-actin rings and fibers. Each resolution step in the contracting path is composed of two sets of 3 × 3 convolutional layers, followed by a batch normalization and a 2 × 2 max-pooling. The layer sizes are of {16, 32, 64, 128} filters respectively. The layers in the expanding path are symmetrical to the contracting path, but with an additional 2 × 2 transposed convolution (stride of 2). As in the seminal implementation, skip links allow to keep and propagate information at various scales. A final 1×1 convolutional layer collapses the features into two segmentation maps, one for the F-actin rings and one for the F-actin fibers (Fig. 4a). Since a region can contain both rings and fibers, we treated each network output independently. Rectified linear unit (ReLU) activation was used throughout the network, except for the output layer, which uses a sigmoid.

The loss function was a root mean squared error (RMSE), and we used early stopping on the validation set to ultimately keep a model with good generalization properties. The Adam optimizer was used with a learning rate of 0.001, with the other parameters kept to their default values. Batch size was set at 72.

To obtain a binary segmentation map, a hard threshold had to be applied on the network predictions. To compute the optimal thresholds for the two independent channels, we generated receiver operating curves (ROC) using the validation set. We computed the euclidean distance between the false positive and true positive rate (FPR, TPR) coordinate of the thresholds to the optimal operation point (FPR=0, TPR=1) and took the threshold having the smallest distance (Supplementary Fig. 7). The optimal threshold (rings: 0.25, fibers: 0.4) was obtained from the median of the thresholds calculated on the ROC curves of 15 low activity and 8 high activity images. Note that these thresholds could be easily adjusted to adapt to different imaging conditions in other experiments.

We trained a total of 25 networks with the same configuration and asserted their performance using common metrics (see section Performance Metrics, Supplementary Fig. 7d). To control for the ability of h_d_ to robustly detect the reorganization of the F-actin patterns in dendrites, we compared the proportions of rings and fibers predicted by the FCN h_d_ or labeled by an expert. Supplementary Fig. 7c shows no significant difference between expert and FCN labeling for both low and high neuronal activity conditions, indicating that the deep learning approach can reliably detect different proportions of F-actin rings and fibers in neurons. We also performed a dimensionality reduction on the embedded central layer of the FCN h_d_ using the non-linear approach Uniform Manifold Approximation and Projection (UMAP) [41]. Supplementary Fig. 15 shows a clear separation in this reduced space between crops containing rings and fibers.

The h_a_ network was designed to detect the presence of the axonal F-actin rings. The resolution steps were the same as for h_d_, but the layer size were of {8, 16, 32, 64} filters respectively (Supplementary Fig. 6a). The layers in the expanding path are symmetrical to the contracting path. The Adam optimizer was used with similar parameters as h_d_, with early stopping used here as well. Batch size was set at 96. Cross entropy was used as a loss function. The binary segmentation map was obtained using a hard threshold of 0.02 on the prediction of the network. This threshold was calculated using the same ROC procedure as for the h_d_ (Supplementary Fig. 6c). We evaluated the metrics on the testing dataset to obtain the performance of the network in generalization (Supplementary Fig. 6d). Examples of predictions from the testing dataset compared to the labels of the expert and the confusion matrix demonstrate the performance of h_a_ (Supplementary Fig. 6e, f).

To control for the sensitivity of h_a_ to detect changes of the axonal F-actin pattern, we tested its capacity to detect the disruption of the periodical lattice on a sample treated with Latrunculin A (Lat A). Supplementary Fig. 16 shows that the deep learning approach can successfully identify an significant reduction of the detected axonal F-actin rings on the Lat A treated cells (Supplementary Fig. 16, 5 neurons, one-sided ANOVA, *p* = 0.049).

### Weak Labels

Due to the time consuming and difficult task related to precise segmentation of numerous super-resolution microscopy images, we formulated the segmentation problem as a weakly supervised learning approach, where the network is trained on coarse labels that are faster and easier to produce. Our results demonstrate the ability of the network to go beyond the training labels and exceed the accuracy of the bounding boxes (B). In other words, the network is able to infer precise segmentation rules using only coarse examples. To assess the improvement in segmentation performance, we asked the expert to take the time to precisely label (P) 70 regions (38 for rings and 32 for fibers) randomly sampled from the testing dataset. We then compared the precision-recall curve (PR), an analysis method that is insensitive to class imbalance (e.g. number of background pixels in our images being more important than the number of foreground pixels) (Fig. 5b). The comparison of the predicted segmentation (S) and B with P as ground truths demonstrated how S exhibits a higher precision over all range of recall than B, showing that the prediction of h_d_ is closer to P than B (Fig. 5b and Supplementary Fig. 9d). This asserts that the trained FCN generated more accurate segmentation than the labels it was trained with. If the network had not gone further than the bounding box labels, its performance on precise labels would have been at most the performance of B on P.

We also set out to analyse how coarse labels can impact on the precision of the predictions. To do so, we trained 5 different instances of networks for different stepwise dilation of the labels (from 100 nm to 1 *µ*m, Fig. 9c,d). Following training, the thresholds for the prediction of rings and fibers were calculated using the ROC procedure described in the Fully Convolutional Neural Network Methods section. The precise dataset was then used with all instances of network to calculate the average precision score (AP) with the precise labels as ground truth (Fig. 5d, Supplementary Fig. 9d).

### Resampling

For bivariate data analysis two resampling methods were employed: 1) bootstrapping [42] for data visualization and 2) randomization test (shuffling) for statistical analysis [43]. The bootstrapping experiments consisted in sampling the data points for each condition with replacement to generate an estimate of the mean values distribution. For each condition (N data points per condition), a bootstrap sample consisted of randomly selecting N data points with replacement from the raw data and calculating the mean value of the distribution. The number of repetitions was set to 10 000 for all conditions. It resulted in a distribution of the bootstrapped mean values that could be used for data visualization (Fig. 6c and 7a,c). For the comparison between two different stimulation conditions, statistical analysis was performed using a randomization test. The null hypothesis for each set of two conditions (A and B) was that both conditions belonged to the same distribution. First, the difference between the mean values of A and B was calculated (*D*_*raw*_ = *µ*_*A*_ − *µ*_*B*_). Data points from both conditions were randomly reassigned to two groups A’ and B’ having *N*_*A*_ and *N*_*B*_ data points respectively. The difference between the mean values of A’ and B’ was determined (*D*_*rand*_ = *µ*_*A*_′ − *µ*_*B*_′). The difference of mean values obtained from the raw data (*D*_*raw*_) was compared to the distribution of the differences obtained from 10 000 randomization samples (*D*_*rand*_) to verify the null hypothesis, assuming multivariate normal distribution of randomization samples.

### Performance Metrics

We define common pixel-wise metrics that were used to evaluate the performance of the architectures [44]. The first metric is the F1-score and measures the overlap between the segmentation map and the ground truth. The F1-score is defined as

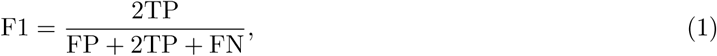

where TP, FP and FN are true-positive, false-positive and false-negative respectively. The second metric calculates the proportion of the ground truth also present in the predicted segmentation. It is called the sensitivity and is defined as

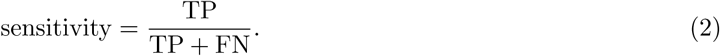

The specificity measures the proportion of background also predicted as background in the segmentation map. It is defined as

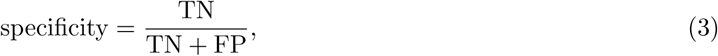

where TN stands for true-negative. Precision, which measures the proportion of true features over all the true prediction, is defined as

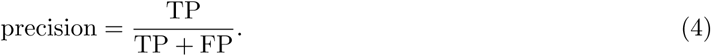

### Analysis of axonal and dendritic branch lengths

For the morphological analysis of the dendritic and axonal processes we used two measurements that were performed on each image: i) total length of analysed branches and ii) maximal length of a branch. Briefly, we generated a skeleton image from the binary masks used to identify dendrites and axons. We extracted all nodes, *i.e.* a joint or an end point, from the skeleton image. The skeleton was subdivided in multiple segments, where each segment has a start and an end node. The total length of branches per image was calculated using the sum of each segment length in the skeleton image. To calculate the maximal branch length per image, we used the NetworkX python library to efficiently create and manipulate a complex network. The complex network (or graph) was created from an adjacency matrix. The weights of the adjacency matrix were set according to the length of the segment between two connected nodes. The weight is set to 0 if two nodes are not connected by a segment.

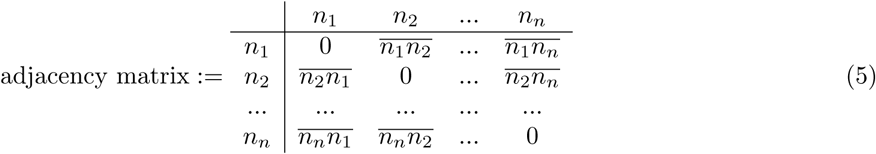

The 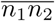 notation represents the length of the segment between nodes *n*_1_ and *n*_2_. While we could have reported the maximal branch length (multiple connected segments) in the graph, we found that in some cases it produced branches that were not representative of the analysed neuronal processes. For this reason, we extracted the shortest path between the two most distant and connected nodes in the graph. This method provided more linear branches and allowed us to report the maximal length of a branch for each image.

## Data and code availability

The datasets for the training of both networks and the dataset for the analysis are available from the corresponding author upon request.

Open source code for the segmentation of the periodical lattice in dendrite and axon is available online: https://github.com/FLClab/STEDActinFCN.

## Acknowledgments

Laurence Emond for sample preparation and immunocytochemistry. Francine Nault, Charleen Salesse and Laurence Emond for the neuronal cell culture. Jonathan Marek and Renaud Bernatchez for the development of a custom Python annotation application. Vincent Poiré and Azadeh Sadat Mozafari for preliminary experiments with U-Net implementation and Gabriel Leclerc for assistance with the skeleton analysis. Armen Saghatelyan for comments on the manuscript. Funding was provided by grants from the Natural Science and Engineering Research Council of Canada (P.D.K., C.G.), the Canadian Institute of Health Research (P.D.K.), MITACS (C.G.), and the Canadian Foundation for Innovation (P.D.K.).

## Author Contributions

F.L.C., M.L. and G.L. performed STED imaging, T.W, F.L.C. and G.L. performed live-cell STED imaging, F.L.C., M.L., G.L. and T.W. prepared samples, F.L.C. generated the labeled datasets, A.B., M.A.G. and C.G. designed the FCN, A.B. implemented the FCN based analysis, F.L.C., A.B. and M.A.G. analyzed the results, F.L.C., M.L. and P.D.K. designed the experiments, F.L.C., M.L., A.B. and P.D.K. wrote the manuscript, F.L.C., C.G. and P.D.K. supervised the project.

## Additional Information

The authors declare no financial and non-financial competing interests.

## References

[1] Xu, K., Zhong, G. & Zhuang, X. Actin, spectrin, and associated proteins form a periodic cytoskeletal structure in axons. Science 339, 452–456 (2013).

[2] D’Este, E., Kamin, D., Göttfert, F., El-Hady, A. & Hell, S. W. STED nanoscopy reveals the ubiquity of subcortical cytoskeleton periodicity in living neurons. Cell Reports 10, 1246–1251 (2015).

[3] Zhong, G. et al. Developmental mechanism of the periodic membrane skeleton in axons. eLife 3, e04581 (2014).

[4] Han, B., Zhou, R., Xia, C. & Zhuang, X. Structural organization of the actin-spectrin–based membrane skeleton in dendrites and soma of neurons. Proceedings of the National Academy of Sciences 114, E6678–E6685 (2017).

[5] Ganguly, A. et al. A dynamic formin-dependent deep F-actin network in axons. Journal of Cell Biology 210, 401–417 (2015).

[6] Konietzny, A., Bär, J. & Mikhaylova, M. Dendritic actin cytoskeleton: structure, functions, and regulations. Frontiers in Cellular Neuroscience 11, 147 (2017).

[7] Schätzle, P. et al. Activity-dependent actin remodeling at the base of dendritic spines promotes microtubule entry. Current Biology 28, 2081–2093 (2018).

[8] Bär, J., Kobler, O., Van Bommel, B. & Mikhaylova, M. Periodic F-actin structures shape the neck of dendritic spines. Scientific Reports 6, 37136 (2016).

[9] Unsain, N. et al. Remodeling of the actin/spectrin membrane-associated periodic skeleton, growth cone collapse and F-actin decrease during axonal degeneration. Scientific Reports 8, 3007 (2018).

[10] Wang, G. et al. Structural plasticity of actin-spectrin membrane skeleton and functional role of actin and spectrin in axon degeneration. eLife 8, e38730 (2019).

[11] Zhou, R., Han, B., Xia, C. & Zhuang, X. Membrane-associated periodic skeleton is a signaling platform for rtk transactivation in neurons. Science 365, 929–934 (2019).

[12] Sidenstein, S. C. et al. Multicolour multilevel STED nanoscopy of actin/spectrin organization at synapses. Scientific Reports 6, 26725 (2016).

[13] Basu, S. & Lamprecht, R. The role of actin cytoskeleton in dendritic spines in the maintenance of long-term memory. Frontiers in Molecular Neuroscience 11, 143 (2018).

[14] Chazeau, A. & Giannone, G. Organization and dynamics of the actin cytoskeleton during dendritic spine morphological remodeling. Cellular and Molecular Life Sciences 73, 3053–3073 (2016).

[15] Ronneberger, O., Fischer, P. & Brox, T. U-net: Convolutional networks for biomedical image segmentation. In International Conference on Medical image computing and computer-assisted intervention, 234–241 (Springer, 2015).

[16] Van Valen, D. A. et al. Deep learning automates the quantitative analysis of individual cells in live-cell imaging experiments. PLoS Computational Biology 12, e1005177 (2016).

[17] Sadanandan, S. K., Ranefall, P., Le Guyader, S. & Wählby, C. Automated training of deep convolutional neural networks for cell segmentation. Scientific Reports 7, 7860 (2017).

[18] Raza, S. E. A. et al. Mimo-net: A multi-input multi-output convolutional neural network for cell segmentation in fluorescence microscopy images. In 2017 IEEE 14th International Symposium on Biomedical Imaging (ISBI 2017), 337–340 (IEEE, 2017).

[19] Falk, T. et al. U-net: deep learning for cell counting, detection, and morphometry. Nature methods 16, 67 (2019).

[20] Papandreou, G., Chen, L.-C., Murphy, K. P. & Yuille, A. L. Weakly-and semi-supervised learning of a deep convolutional network for semantic image segmentation. In Proceedings of the IEEE International Conference on Computer Vision, 1742–1750 (2015).

[21] Akram, S. U., Kannala, J., Eklund, L. & Heikkilä, J. Cell segmentation proposal network for microscopy image analysis. In Deep Learning and Data Labeling for Medical Applications, 21–29 (Springer, 2016).

[22] Kraus, O. Z., Ba, J. L. & Frey, B. J. Classifying and segmenting microscopy images with deep multiple instance learning. Bioinformatics 32, i52–i59 (2016).

[23] De Koninck, P., Carbonetto, S. & Cooper, E. Ngf induces neonatal rat sensory neurons to extend dendrites in culture after removal of satellite cells. Journal of Neuroscience 13, 577–585 (1993).

[24] Urban, N. T., Willig, K. I., Hell, S. W. & Nägerl, U. V. Sted nanoscopy of actin dynamics in synapses deep inside living brain slices. Biophysical journal 101, 1277–1284 (2011).

[25] Frost, N. A., Shroff, H., Kong, H., Betzig, E. & Blanpied, T. A. Single-molecule discrimination of discrete perisynaptic and distributed sites of actin filament assembly within dendritic spines. Neuron 67, 86–99 (2010).

[26] Wegner, W. et al. In vivo mouse and live cell sted microscopy of neuronal actin plasticity using far-red emitting fluorescent proteins. Scientific reports 7, 11781 (2017).

[27] Bubb, M. R., Spector, I., Beyer, B. B. & Fosen, K. M. Effects of jasplakinolide on the kinetics of actin polymerization an explanation for certain in vivo observations. Journal of Biological Chemistry 275, 5163–5170 (2000).

[28] Lukinavičius, G. et al. Fluorogenic probes for live-cell imaging of the cytoskeleton. Nature Methods 11, 731 (2014).

[29] Mortal, S. et al. Actin waves do not boost neurite outgrowth in the early stages of neuron maturation. Frontiers in Cellular Neuroscience 11, 402 (2017).

[30] Lu, W.-Y. et al. Activation of synaptic nmda receptors induces membrane insertion of new ampa receptors and ltp in cultured hippocampal neurons. Neuron 29, 243–254 (2001).

[31] Hudmon, A. et al. A mechanism for Ca^2+^/calmodulin-dependent protein kinase II clustering at synaptic and nonsynaptic sites based on self-association. Journal of Neuroscience 25, 6971–6983 (2005).

[32] Vohs, K. D. et al. Making choices impairs subsequent self-control: A limited-resource account of decision making, self-regulation, and active initiative. Journal of Personality and Social Psychology 94, 883–898 (2008).

[33] Umemiya, M., Senda, M. & Murphy, T. H. Behaviour of NMDA and AMPA receptor-mediated miniature EPSCs at rat cortical neuron synapses identified by calcium imaging. The Journal of Physiology 521, 113–122 (1999).

[34] Pollard, T. D. & Cooper, J. A. Actin and actin-binding proteins. a critical evaluation of mechanisms and functions. Annual Review of Biochemistry 55, 987–1035 (1986).

[35] Lynch, G. & Baudry, M. Brain spectrin, calpain and long-term changes in synaptic efficacy. Brain Research Bulletin 18, 809–815 (1987).

[36] Albrecht, D. et al. Nanoscopic compartmentalization of membrane protein motion at the axon initial segment. Journal of Cell Biology 37–46 (2016).

[37] Wang, Z. et al. Myosin Vb mobilizes recycling endosomes and ampa receptors for postsynaptic plasticity. Cell 135, 535–548 (2008).

[38] Durand, A. et al. A machine learning approach for online automated optimization of super-resolution optical microscopy. Nature communications 9 (2018).

[39] Schindelin, J. et al. Fiji: an open-source platform for biological-image analysis. Nature methods 9, 676–682 (2012).

[40] Otsu, N. A threshold selection method from gray-level histograms. IEEE Transactions on Systems, Man, and Cybernetics 9, 62–66 (1979).

[41] McInnes, L. & Healy, J. Umap: Uniform manifold approximation and projection for dimension reduction. arXiv preprint 1802.03426 (2018).

[42] Efron, B. & Tibshirani, R. J. An introduction to the bootstrap (CRC press, 1994).

[43] Good, P. I. Resampling Methods (Birkhäuser Basel, 2006), 3 edn.

[44] Yeghiazaryan, V. & Voiculescu, I. An overview of current evaluation methods used in medical image segmentation. Department of Computer Science, University of Oxford (2015).

