## Supplemental Figures for "Neuronal activity remodels the F-actin based submembrane lattice in dendrites but not axons of hippocampal neurons"

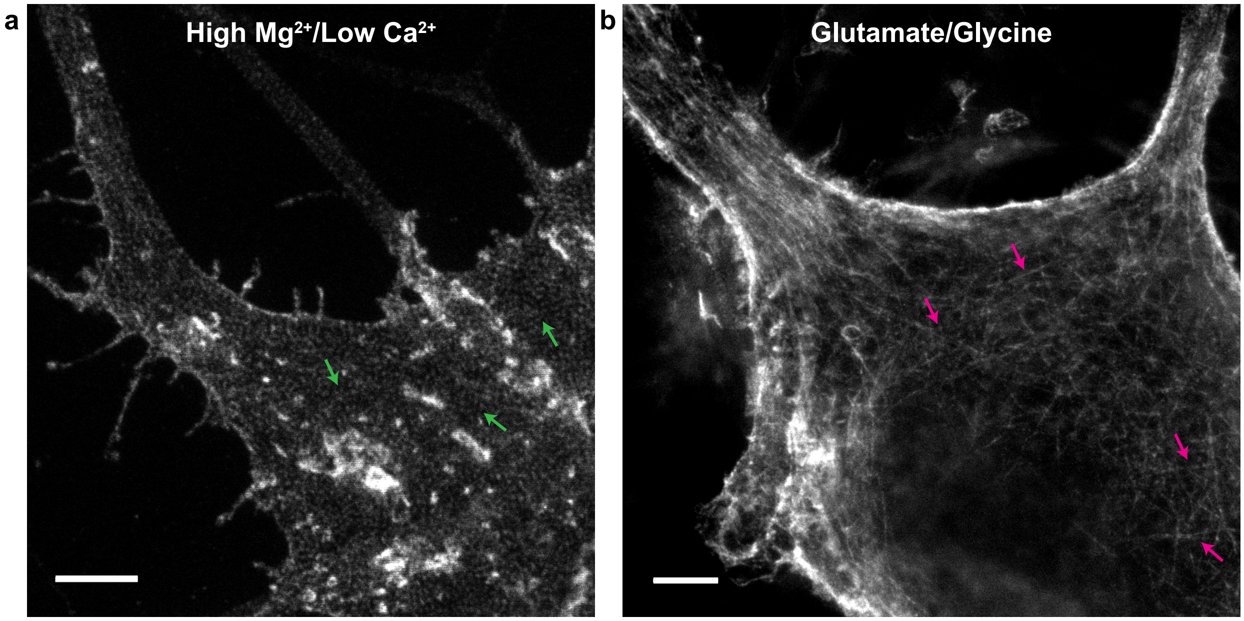


Supplementary Figure 1: Regions of the soma showing a) the 2D polygonal F-actin lattice structure (green arrows) [1] for the low activity High $Mg^{2+}$/low $Ca^{2+}$ condition and b) the thin actin fibers (magenta arrows) with the Glutamate/Glycine stimulation condition. Scale bars 2$\mu$m

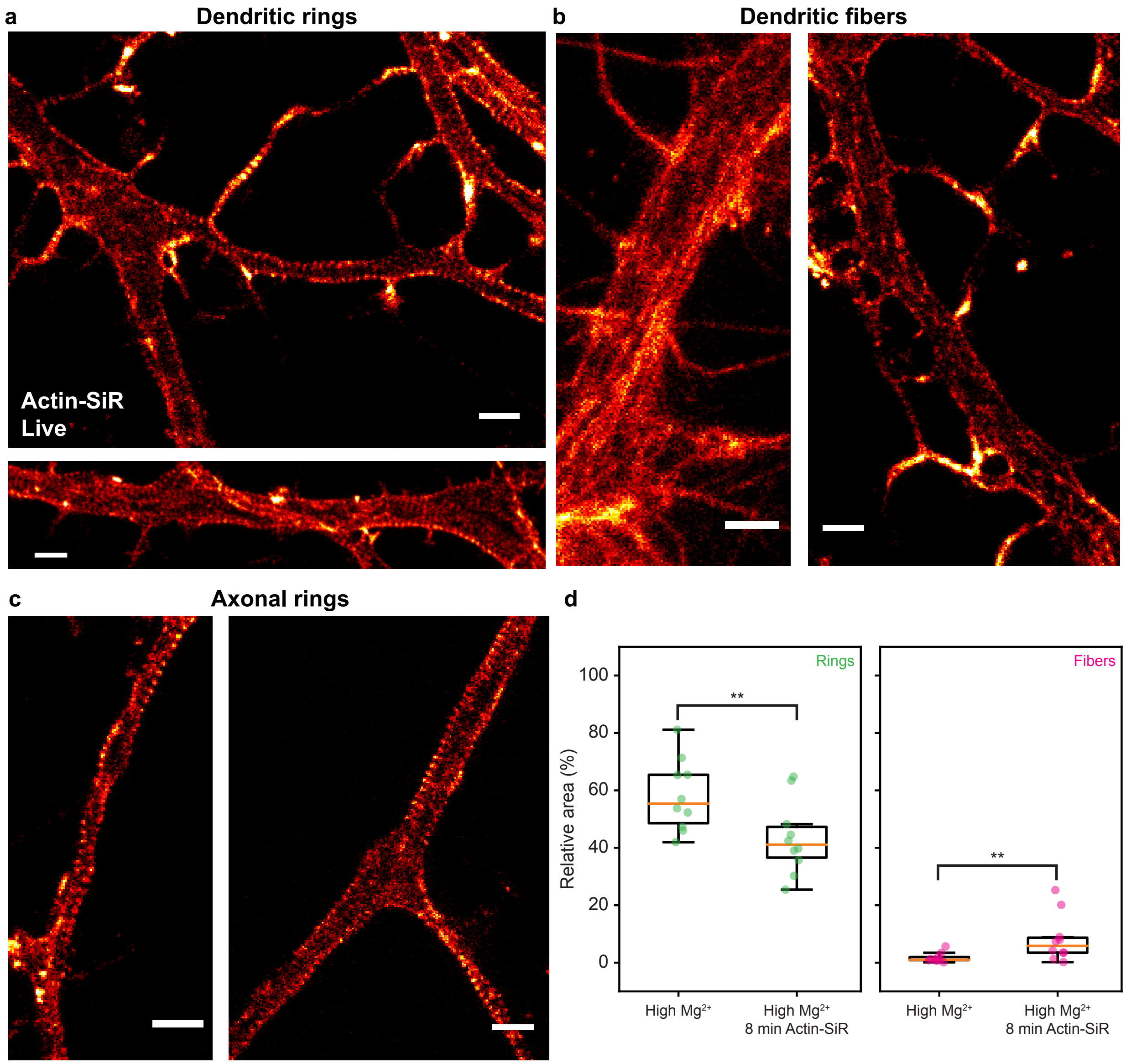


Supplementary Figure 2: Staining of the F-Actin lattice with Actin-SiR. Live-cell STED imaging of the F-actin patterns in dendrites and axons. Representative images of live hippocampal neurons stained with Actin-SiR (8 min in 0.5 $\mu$M) showing a) the dendritic F-actin rings, b) the longitudinal fibers in dendrites, and c) the axonal rings. Live-cell imaging of neurons stained with Actin-SiR was performed on more than 20 neurons from 5 different cultures d) Incubation of live hippocampal neurons with Actin-SiR significantly reduces the detected F-actin ring pattern and increases the detected fibers in dendrites compared to controls ($p_{\text{Rings}} = 0.005$ and $p_{\text{Fibers}} = 0.008$, N = 10 for each condition). Actin-SiR (0.5 $\mu$M) was incubated for 8 min in the growth media. Subsequently, the neurons were transferred in High $Mg^{2+}$/low $Ca^{2+}$ blocking solution for 10 min and fixed immediately in 4% PFA solution (See Methods). The area of dendritic rings and fibers was analysed using the FCN $h_d$ for the control and the neurons that were pre-treated with Actin-SiR. Scale bars 2 $\mu$m.

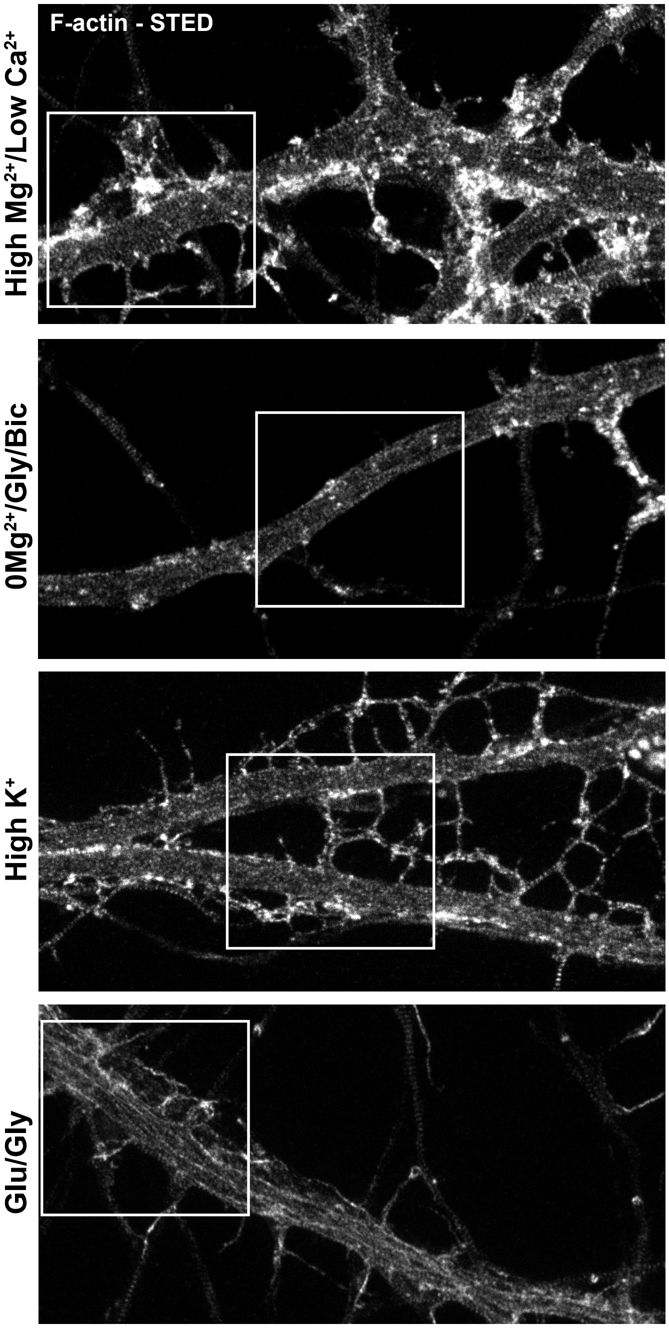


Supplementary Figure 3: Raw F-actin signal without overlay from the images shown in Fig. 2.

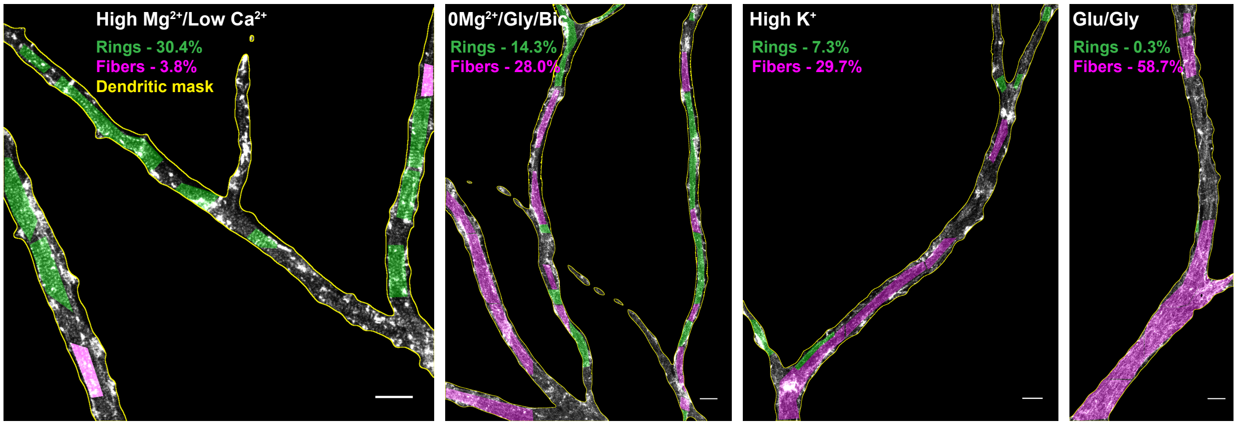


Supplementary Figure 4: Representative images manually labelled with bounding boxes to measure the relative areas of F-actin rings (green) and longitudinal fibers (magenta) within the dendritic mask (yellow line). Those images are part of the FCN training dataset. Scale bars 1 $\mu$m.

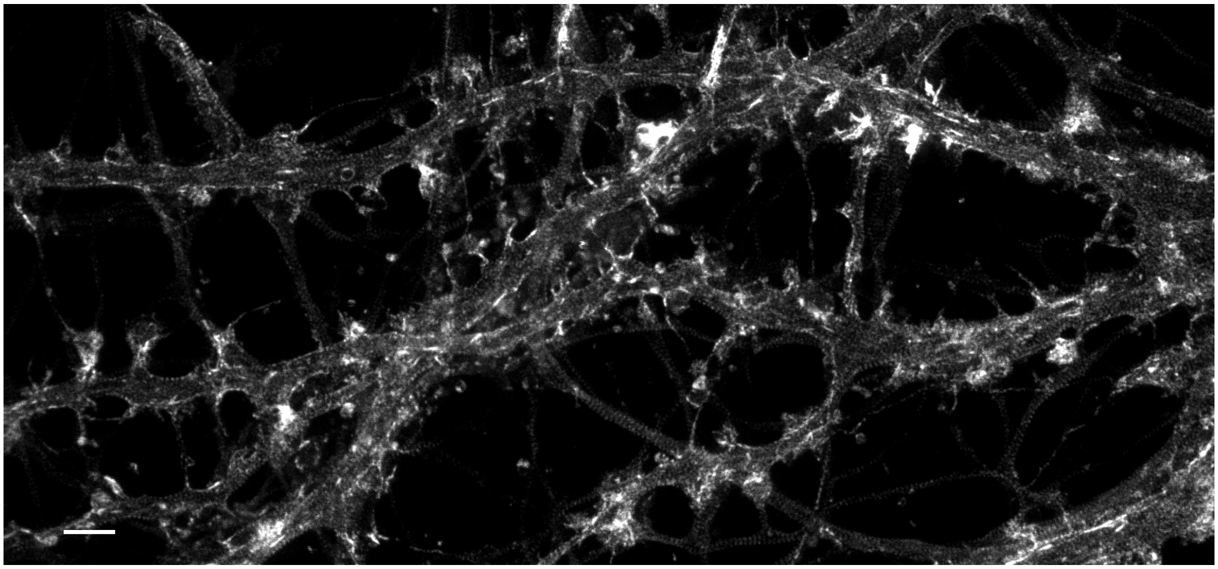

Supplementary Figure 5: Raw F-actin signal without overlay from the images shown in Fig. 4. Scale bar 2 $\mu$m.

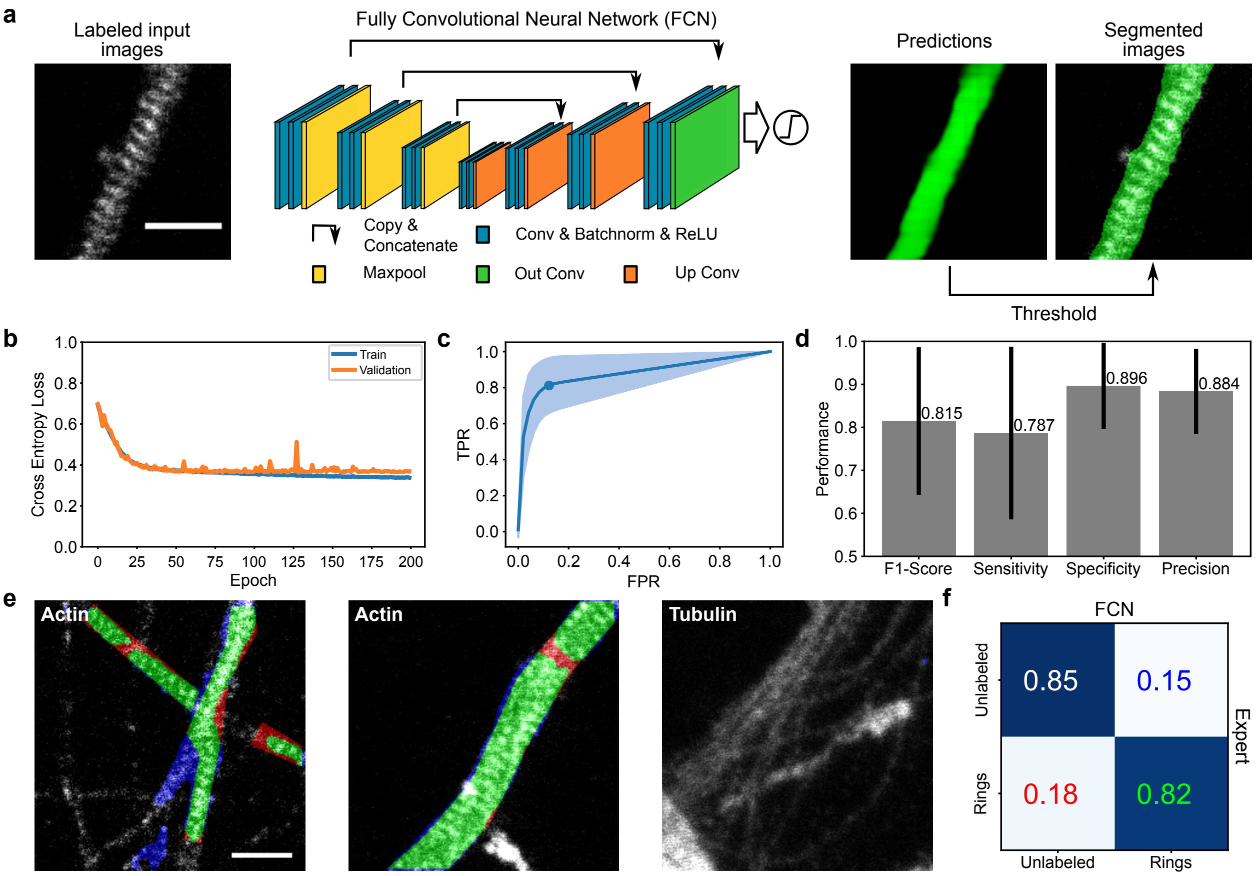


Supplementary Figure 6: Performance evaluation of the FCN $h_a$. a) A very similar architecture to $h_d$ was chosen for $h_a$ (see Materials and Methods for specific implementation details, Conv : Convolution, Batchnorm : Batch Normalization, ReLU : Rectified Linear Unit). Since no fibers were detected in axons by the expert, the segmentation task was simplified to a single class (F-actin rings) for $h_a$. The network takes as an input an image and outputs a prediction (in range $[0, 1]$) for the presence of the F-actin ring pattern. b) Learning curves showing the evolution of the objective function during training (blue) and validation (orange). The average (solid line) and standard deviation (pale region) of the loss was calculated at every epoch. c) Mean Receiver Operating Characteristic (ROC, FPR : false positive rate, TPR : true positive rate) curve (blue line, STD: pale blue) calculated on the validation dataset (112 images) was used to determine the optimal segmentation threshold (0.02). d) Performance of the FCN $h_a$ in the segmentation of F-actin rings regarding the F1-score, sensitivity, specificity and precision on all images of the testing dataset (69 images, See Methods). A metric performance of 70% was reported to be a good segmentation by Falk *et al* [2], while 90% is close to human annotations. Performance was averaged from all images (black line: STD). e) Examples of axonal segments stained for $\alpha$-tubulin or F-actin from the testing dataset. As expected, no axonal F-actin rings are predicted by the FCN on $\alpha$-tubulin images. True positives are shown in green, false positive in blue, and false negative in red. f) The confusion matrix of the network $h_a$ calculated on all images from the testing dataset. The numbers are colored according to the color-code of the segmentation map shown in e. True positives are shown in green (rings) and magenta (fibers), false positive in blue, and false negative in red. Scale bars 1 $\mu$m.

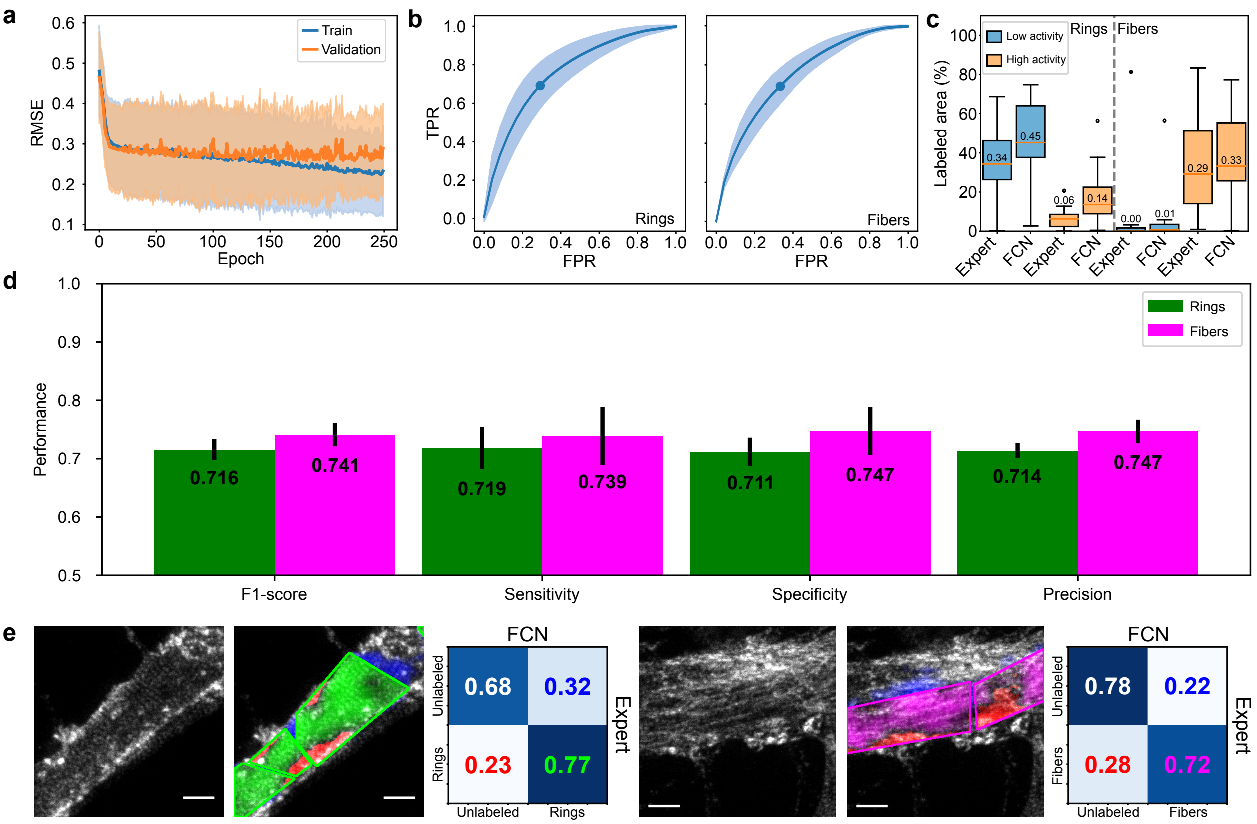


Supplementary Figure 7: Performance evaluation of $h_d$ a) Learning curves of $h_d$ showing the evolution of the objective function during training (blue) and validation (orange). At each epoch and for every training instance (2263 crops), the mean (solid line) and standard deviation (pale region) was computed. b) Mean Receiver Operating Characteristic (ROC, FPR : false positive rate, TPR : true positive rate) curves (blue line) and standard deviation (STD) of the curve (pale blue), obtained from the ROC curves of 15 low activity images (high proportion of F-actin ring pattern) and 8 high activity images (high proportion of longitudinal fibers) from the validation dataset. Using the ROC curves for both structures, the optimal threshold (rings: 0.25, fibers 0.4) was calculated to generate the segmentation maps. c) To characterize the performance of the FCN, we compared the area where F-actin rings (left) and fibers (right) were detected by an expert and the FCN $h_d$ at low (blue) and high (orange) neuronal activity. The dendritic area was determined for each image separately using the dendritic mask from MAP2 staining (see Methods). $n_{\text{low activity}} = 12$, $n_{\text{high activity}} = 14$. d) Performance of the FCN $h_d$ in the segmentation of F-actin rings and fibers regarding the F1-score, sensitivity, specificity and precision on all images of the testing dataset (105 images, See Methods). Performance was averaged from 25 different instances of the same architecture (black line: STD, green: rings, magenta: fibers). e) Performance evaluation using confusion matrices for the FCN-based segmentation of rings (left) and fibers (right) on the complete testing dataset (105 images). True positives are shown in green (rings) and magenta (fibers), false positive in blue, and false negative in red. Scale bars 1 $\mu$m.

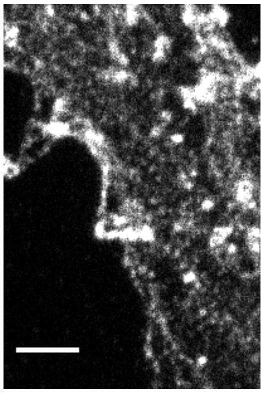

Supplementary Figure 8: Raw F-actin signal without overlay from the image shown in Fig. 5a. Scale bar 1 $\mu$m.

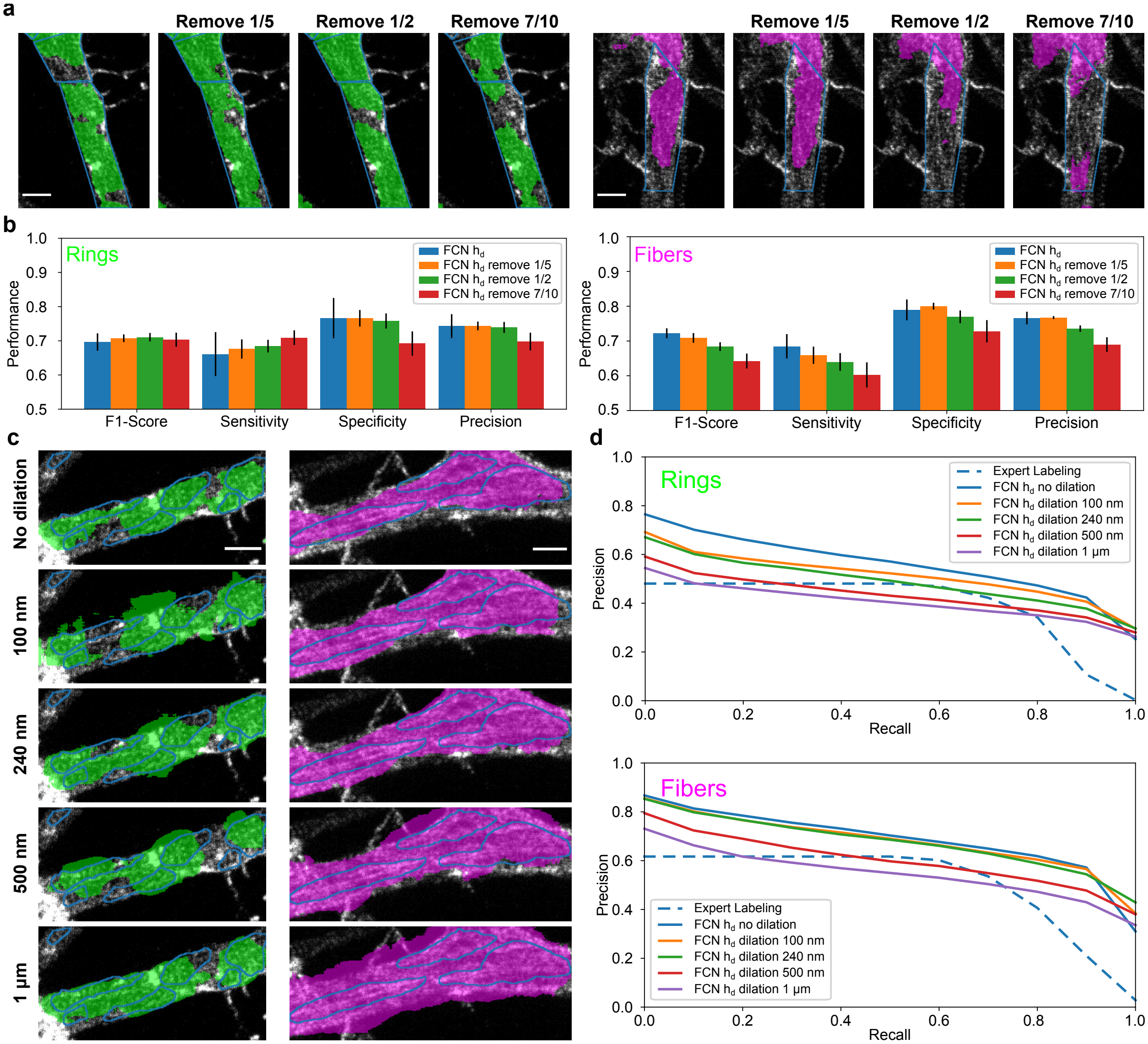


Supplementary Figure 9: Evaluation of the impact of labeling omission and reduced training set labeling precision on the performance of the FCN $h_d$ for the segmentation of dendritic F-actin rings and longitudinal fibers. a) Prediction of $h_d$ on the testing dataset for the presence of F-actin rings (green) or longitudinal fibers (magenta). The corresponding expert bounding box labels are shown in blue. Shown are the results obtained with $h_d$ using training datasets with 4/5, 1/2 and 3/10 of the expert labels but keeping the same number of training crops. b) The performance metrics were evaluated for 5 instances of the network for each training scheme. Black lines represent standard deviation for both graphs. The prediction in a) are shown for a random instance of each network. c) Representative examples of FCN predictions for F-actin rings (green) and longitudinal fibers (magenta) for the networks trained with variable degree of labeling precision. Precise contour labels of the expert are shown in blue. Stepwise uniform dilation of the labels from 100 nm to 1 $\mu$m was used to generate the training datasets. d) The Precision-Recall curves (PR) of 5 different network instances were evaluated for each dilation on the precisely labeled dataset. The mean average precision score (mAP) calculated from the area under the PR curves reported in Fig. 3d show that a dilation of less than 500 nm had better performance than the expert using bounding box annotation for F-actin rings. As for the longitudinal fibers, a dilation of 1$\mu$m still produced more precise labels than the expert on the precisely labeled dataset. Scale bars 1$\mu$m.

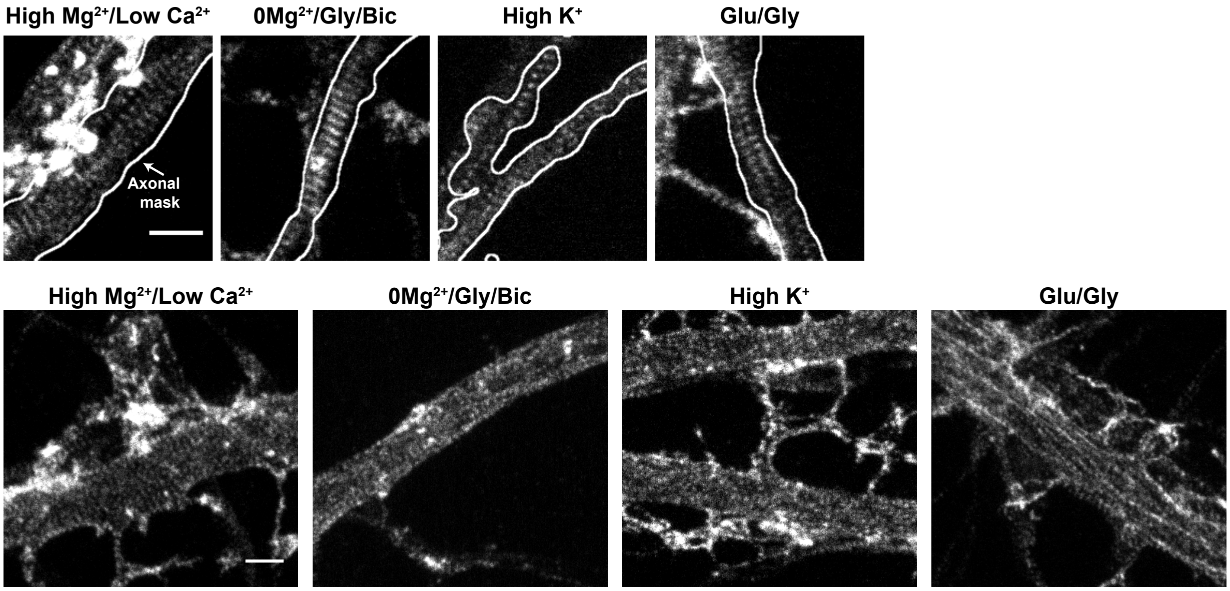


Supplementary Figure 10: Raw F-actin signal without overlay from the image shown in Fig. 6. Scale bars 1 $\mu$m.

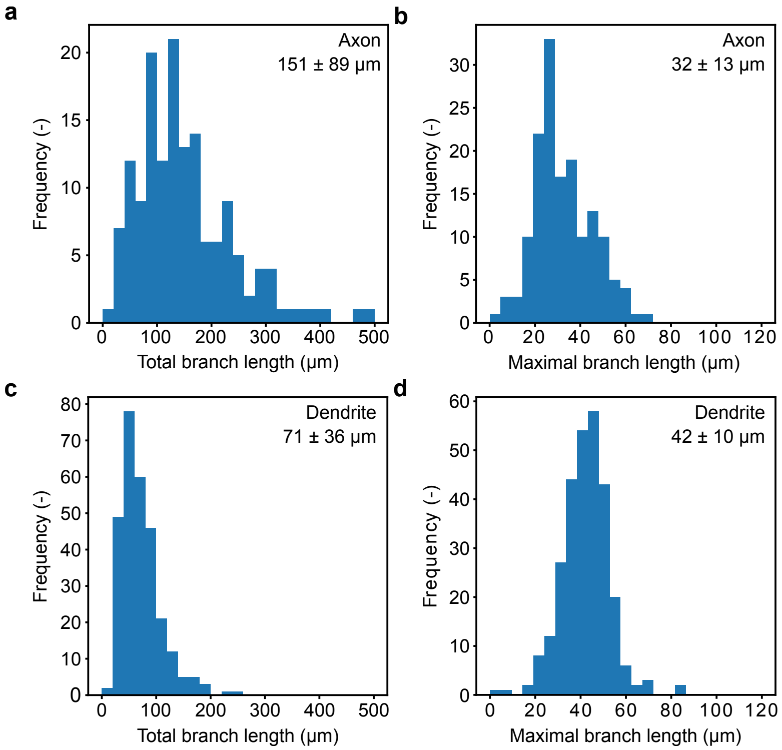


Supplementary Figure 11: Morphological analysis of dendritic and axonal branches length. a,b) A mean total length per image of $151 \pm 89 \mu m$ (a, mean $\pm$ std) and a maximal branch length of $32 \pm 13 \mu m$ (b) were calculated for the analysed axons. c,d). A mean total length per image of $71 \pm 36 \mu m$ (c) and a maximal branch length of $42 \pm 10 \mu m$ (b) are calculated for the analysed dendrites. $N_{\text{axon}} = 150$, $N_{\text{dendrite}} = 287$.See Materials and Methods section for detailed calculation.

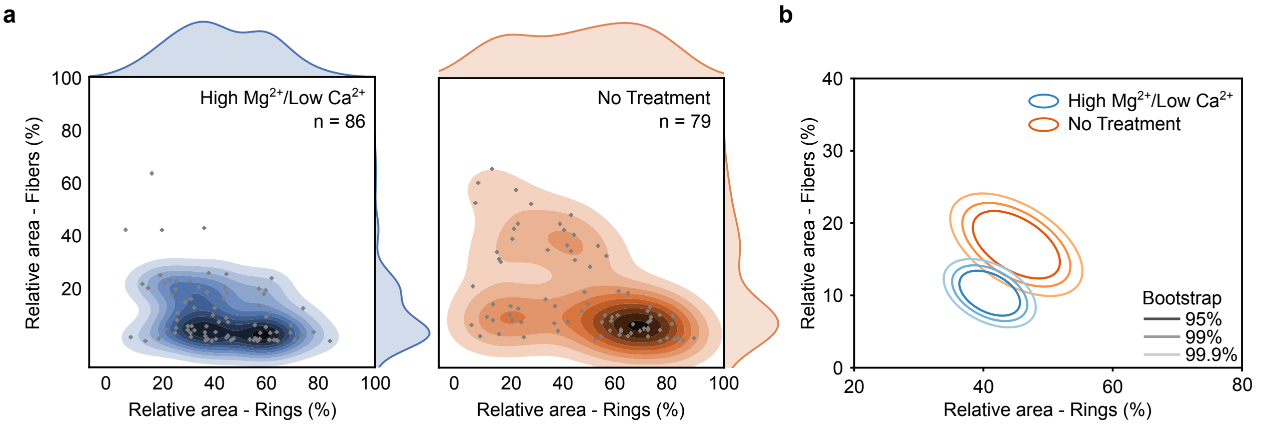


Supplementary Figure 12: Lowering neuronal activity with a High $Mg^{2+}$/Low $Ca^{2+}$ treatment reduces the proportion of F-actin fibers and the variability between neurons. a) Bivariate kernel density estimate of the raw data (grey cross) for 13 DIV neurons treated with i) High $Mg^{2+}$/Low $Ca^{2+}$ (blue) or ii) untreated (orange). b) Mean distributions using bootstrapping. The proportion of F-actin fibers and rings is more variable between different neurons for the basal condition compared to High $Mg^{2+}$/Low $Ca^{2+}$. Due to basal activity, the detected area for F-actin fibers is higher for the untreated condition compared to the High $Mg^{2+}$/Low $Ca^{2+}$. Shown are the regions comprising 95%, 99% and 99.9% of the data point distribution.

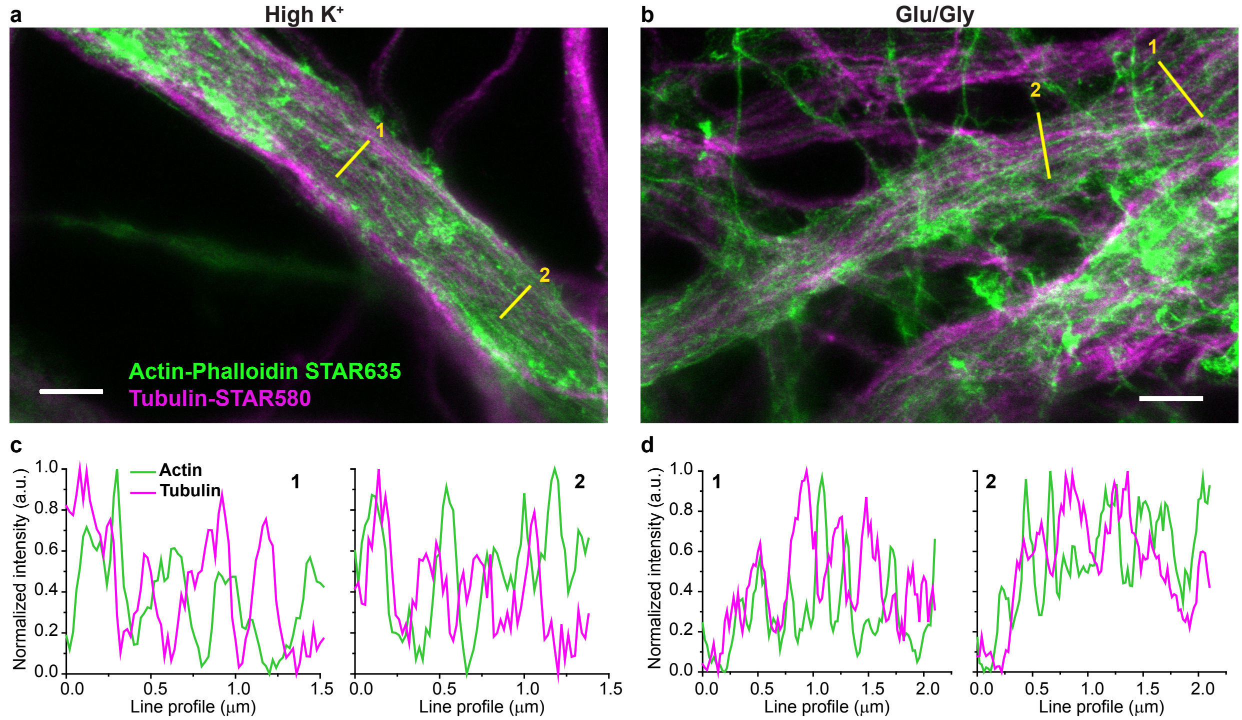


Supplementary Figure 13: F-actin fibers do not colocalize with $\alpha$-tubulin in dendrites. Immunostaining of the two cytoskeleton proteins $\alpha$-tubulin (magenta) and F-actin (green) imaged with dual-color STED nanoscopy showing that the longitudinal actin fibers observed following high $K^+$ (a) or Glutamate/Glycine (b) stimulations do not colocalize with tubulin fibers in dendrites (c&d). Scale bars $2\mu m$

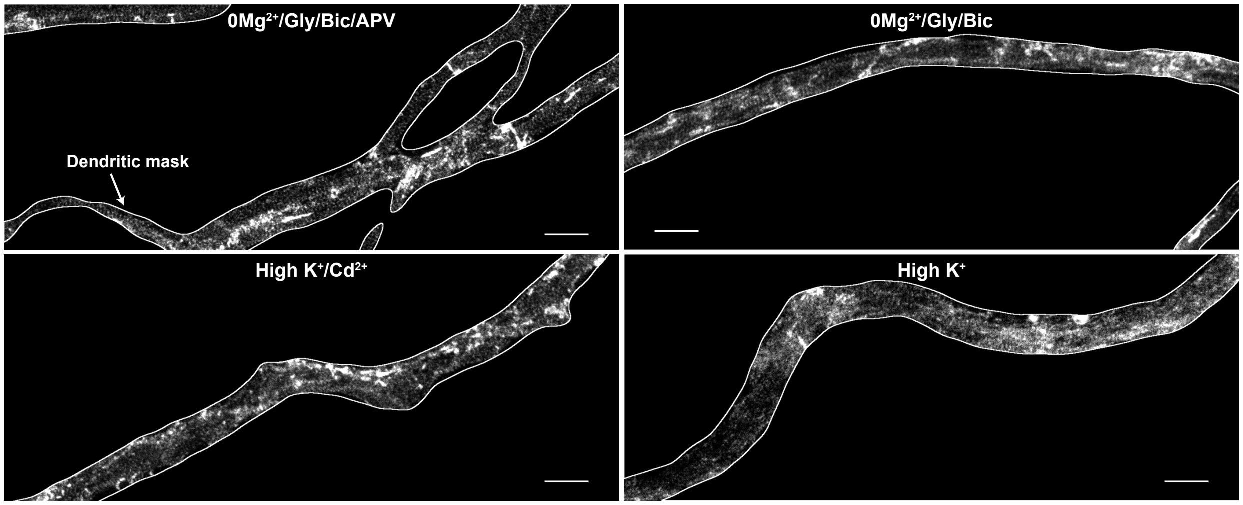


Supplementary Figure 14: Raw F-actin signal without overlay from the image shown in Fig. 6. Scale bars 2 $\mu$m.

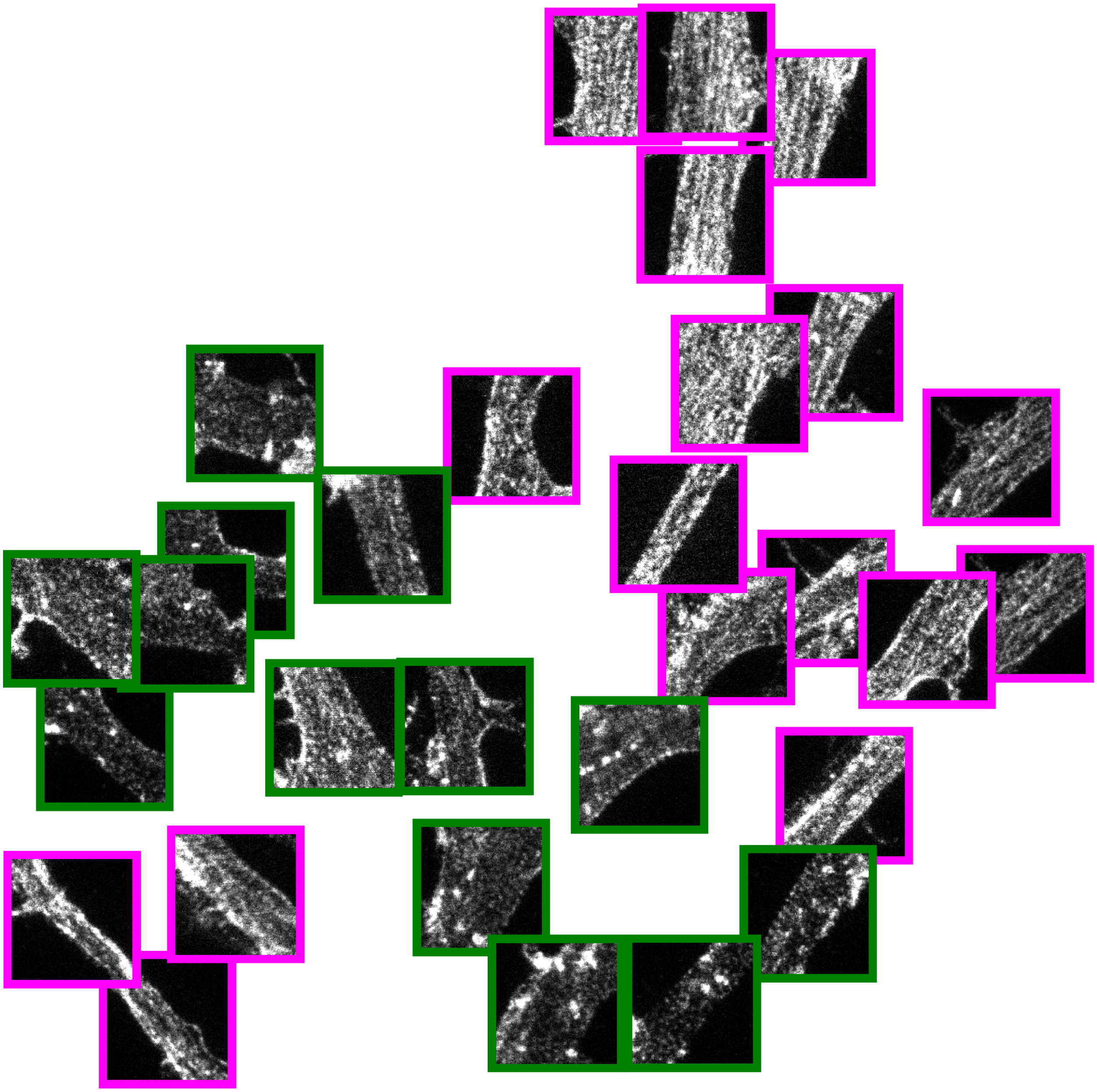

Supplementary Figure 15: UMAP dimension reduction of the central network layer. We performed a UMAP dimension reduction [3] from $[128 \times 16 \times 16]$ of the central FCN $h_d$ layer to 2 dimensions for 30 representative crops from the testing dataset. The border of the images represent the structure that was predicted by the network for each crop (rings : $n = 13$ and fibers $n = 17$). Using this representation, we can clearly distinguish images with rings (green border) and fibers (magenta border) in different region of the space. We also observe that the spatial orientation of the dendritic shaft contributes to the disposition of the images in the UMAP representation.

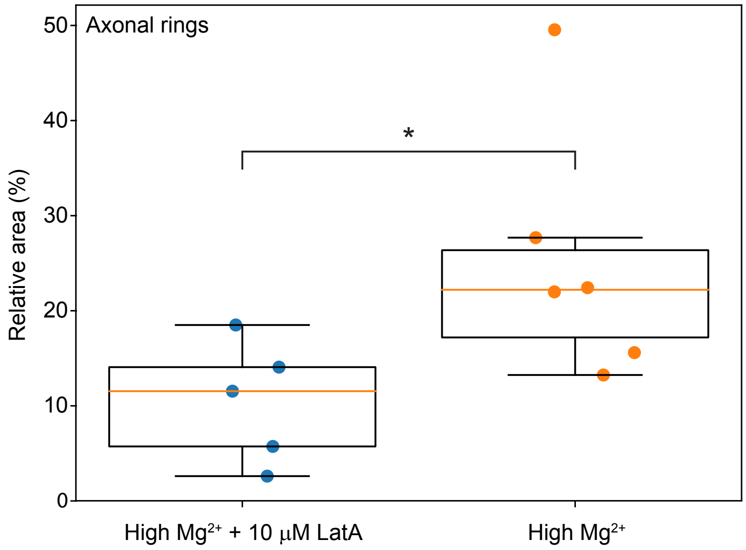


Supplementary Figure 16: Evaluation of the deep learning approach to quantify the F-actin ring disruption in axons. To characterize the ability of the FCN $h_a$ to detect changes in the axonal periodical F-actin rings, cells were treated with Latrunculin A, which has been shown to disrupt this structure in axons [1]. Neurons bathing in High $Mg^{2+}$ blocking solution were treated or not with 10 $\mu$M LatA for 1h. A significant decrease in the axonal area exhibiting periodical F-actin rings was detectable using the FCN $h_a$. $n_{LatA} = 5$, $n_{control} = 6$, $p = 0.049$.

### References

[1] Han, B., Zhou, R., Xia, C. & Zhuang, X. Structural organization of the actin-spectrin–based membrane skeleton in dendrites and soma of neurons. *Proceedings of the National Academy of Sciences* **114**, E6678–E6685 (2017).

[2] Falk, T. *et al.* U-net: deep learning for cell counting, detection, and morphometry. *Nature methods* **16**, 67 (2019).

[3] McInnes, L. & Healy, J. Umap: Uniform manifold approximation and projection for dimension reduction. *arXiv preprint arXiv:1802.03426* (2018).
